# Type III CRISPR–Cas provides resistance against nucleus-forming jumbo phages via abortive infection

**DOI:** 10.1101/2022.06.20.496707

**Authors:** David Mayo-Muñoz, Leah M. Smith, Carmela Garcia-Doval, Lucia M. Malone, Kate R. Harding, Simon A. Jackson, Hannah G. Hampton, Robert D. Fagerlund, Laura F. Gumy, Peter C. Fineran

**Affiliations:** Department of Microbiology and Immunology, University of Otago, PO Box 56, Dunedin 9054, New Zealand; Genetics Otago, University of Otago, PO Box 56, Dunedin 9054, New Zealand; Department of Biochemistry, University of Zürich, CH-8057 Zürich, Switzerland; Bioprotection Aotearoa, University of Otago, PO Box 56, Dunedin 9054, New Zealand; Department of Anatomy, University of Otago, PO Box 56, Dunedin 9054, New Zealand

**Author notes:** Department of Bionanoscience, Delft University of Technology, 2629 HZ Delft, Netherlands. Cawthron Institute, Nelson 7010, New Zealand (Hannah G. Hampton).

## Abstract

Bacteria have diverse defences against phages. In response, jumbo phages evade multiple DNA-targeting defences by protecting their DNA inside a nucleus-like structure. We previously demonstrated that RNA-targeting type III CRISPR–Cas systems provide jumbo phage immunity by recognising viral mRNA exported from the nucleus for translation. Here, we demonstrate that recognition of phage mRNA by the type III system activates a cyclic triadenylate-dependent accessory nuclease, NucC. Although unable to access phage DNA in the nucleus, NucC degrades the bacterial chromosome, triggers cell death, and disrupts phage replication and maturation. Hence, type III-mediated jumbo phage immunity occurs via abortive infection, with suppression of the viral epidemic protecting the population. We further show that type III systems targeting jumbo phages have diverse accessory nucleases, including RNases that provide immunity. Our study demonstrates how type III CRISPR–Cas systems overcome the inaccessibility of jumbo phage DNA to provide robust immunity.

## INTRODUCTION

Clustered regularly interspaced short palindromic repeats (CRISPR) and CRISPR-associated (*cas*) genes provide prokaryotes with sequence-specific protection against viruses and other mobile genetic elements (MGEs) (Barrangou and Marraffini, 2014). These systems consist of (1) CRISPR array(s) of short repeats separated by variable ‘spacer’ sequences complementary to MGEs, and (2) *cas* genes encoding the proteins for immunity. Briefly, mature CRISPR RNAs (crRNAs) generated from CRISPR arrays form interference complexes with Cas proteins and guide them to complementary invader sequences to elicit degradation (Hille et al., 2018; Jackson et al., 2017; Mohanraju et al., 2016). CRISPR–Cas systems are divided into two major classes, six types and multiple subtypes (Makarova et al., 2020b).

Bacterial viruses (bacteriophages or phages) have multiple strategies to counteract CRISPR– Cas and other anti-phage systems (Hampton et al., 2020; Malone et al., 2021; Stanley and Maxwell, 2018). Recently, nucleus-forming jumbo phages that infect *Pseudomonas* and *Serratia* were shown to evade DNA-targeting immune defences, such as type I or II CRISPR– Cas and restriction–modification systems (Malone et al., 2020; Mendoza et al., 2020). Jumbo phages are abundant, defined as having genomes >200 kb and some form a proteinaceous ‘nucleus’ inside the bacterium that encloses phage DNA for replication and transcription (Chaikeeratisak et al., 2017a; Chaikeeratisak et al., 2017b; Malone *et al*., 2020). During virion assembly, phage DNA is directly loaded into phage capsids docked on the outside of the nucleus, which then dissociate, attach to phage tails and exit the cell upon lysis (Chaikeeratisak et al., 2022; Chaikeeratisak *et al*., 2017a; Chaikeeratisak *et al*., 2017b; Nieweglowska et al., 2022). Therefore, the proteinaceous nucleus provides a physical barrier to exclude bacterial defences from targeting phage DNA (Malone *et al*., 2020; Mendoza *et al*., 2020). Nevertheless, jumbo phage mRNA requires export to the bacterial cytoplasm for translation, and bacteria have exploited this circumstance to inhibit phage infection by employing RNA-targeting CRISPR–Cas systems, such as types III and VI (Malone *et al*., 2020; Mendoza *et al*., 2020). Indeed, previous bioinformatics data provide evidence that type III immunity against nucleus-forming jumbo phages is widespread in diverse classes of proteobacteria (Malone *et al*., 2020).

Typically, upon binding complementary phage RNA, type III CRISPR–Cas interference complexes become activated and initiate a multi-step pathway of anti-phage nuclease activities (Varble and Marraffini, 2019; Zhu et al., 2018): (1) the interference complex cleaves nearby phage ssDNA via the Cas10-HD domain (Elmore et al., 2016; Kazlauskiene et al., 2016; Samai et al., 2015b), (2) the Cas10 palm domain synthesises a mixture of cyclic oligoadenylate (cOA) messengers with different ring sizes that activate separate non-specific accessory nucleases (Grüschow et al., 2021; Kazlauskiene et al., 2017; McMahon et al., 2020; Niewoehner et al., 2017; Røstol et al., 2021; Rouillon et al., 2018), and (3) the specific RNA bound to the interference complex is cleaved by each Cas7 (Csm3/Cmr4) subunit (Samai *et al*., 2015b; Staals et al., 2014). This multi-step pathway provides a powerful response to clear phage infection and remains active until the phage DNA is cleared and no longer transcribes any ‘activating’ target RNA (Jiang et al., 2016; Røstol and Marraffini, 2019; Rouillon *et al*., 2018). Accessory nucleases are only essential when interference complexes become overwhelmed by phage DNA accumulation and induce dormancy to ‘buy time’ to clear the invader DNA (Røstol and Marraffini, 2019; Røstol *et al*., 2021). In some cases, cOAs are also enzymatically degraded as a self-regulation mechanism (Athukoralage et al., 2018; Garcia-Doval et al., 2020; Jia et al., 2019).

*Serratia* sp. ATCC 39006 (hereafter *Serratia*) encodes functional type I-E, I-F and III-A CRISPR–Cas systems (Patterson et al., 2016). Our recent work demonstrated that type III, but not type I CRISPR–Cas, protects from jumbo phage infection (Malone *et al*., 2020). Despite bioinformatic evidence that type III are the predominant CRISPR–Cas systems to provide jumbo phage immunity (Malone *et al*., 2020), phage PCH45–*Serratia* is the only current experimental model of type III immunity against nucleus-forming jumbo phages. Importantly, the mechanism by which type III immunity is elicited against jumbo phages has not been resolved. The type III-A interference complex cannot access jumbo phage DNA in the nucleus, and inactivation of the Cas10-HD DNase has no impact on immunity (Malone *et al*., 2020). Therefore, type III CRISPR–Cas must function through an alternative mechanism to protect from jumbo phages as the conventional type III model requires access to, and degradation of, phage DNA to clear the infection (Jiang *et al*., 2016; Røstol and Marraffini, 2019; Samai et al., 2015a). Interestingly, RNA recognition, the Cas10 palm domain and an accessory putative nuclease were required for jumbo phage defence (Malone *et al*., 2020).

The *Serratia* type III accessory nuclease is related to the recently discovered NucC superfamily of DNA endonucleases associated with CBASS defence systems (Lau et al., 2020; Malone *et al*., 2020; Ye et al., 2020). In some CBASS systems, detection of specific peptides by a HORMA domain protein stimulates cyclic triadenylate (cA3) synthesis, activating non-specific DNase activity of NucC, which triggers abortive infection (Abi) (Ye *et al*., 2020). During Abi, infected cells die before the phage can complete its replication cycle, preventing the phage epidemic from spreading to nearby cells, thus protecting the bacterial population (Lopatina et al., 2020). Some type III CRISPR–Cas systems, including that of *Serratia*, have accessory genes encoding NucC homologues activated by cA3 (Grüschow *et al*., 2021; Lau *et al*., 2020), but their role in phage defence is untested.

Here, we address how type III CRISPR–Cas can provide immunity against jumbo phages that protect their DNA in a nucleus-like structure. Specifically, we demonstrate how the *Serratia* type III-A CRISPR–Cas system provides resistance against the jumbo phage PCH45, including the role of the accessory nuclease NucC and the consequences for individual bacteria and the wider population. We discovered that NucC cannot access phage DNA, but triggers Abi by destruction of the bacterial genome, and while individual infected cells die, the population survives at low phage doses. Importantly, type III-mediated jumbo phage immunity is a widespread mechanism that can be elicited by multiple families of accessory nucleases, including RNases.

## RESULTS

### *Serratia* NucC forms a hexamer that binds cA3

Since resistance against jumbo phage PCH45 required a *Serratia* type III-A accessory gene with homology to the NucC nuclease (Malone *et al*., 2020), we explored its mechanism as part of CRISPR–Cas immunity. *Serratia* NucC contains 250 amino acids (28.14 kDa) and shares <35% sequence identity with recently characterized NucC proteins from CBASS and a type III-B CRISPR–Cas system (Lau *et al*., 2020; Ye *et al*., 2020) (**Figure 1A-B**). Despite the low identity, the active site motif of ID–30ExK in these restriction endonuclease-like fold proteins is conserved in *Serratia* NucC (**Figure 1B**), suggesting it may also function as an endonuclease. To elucidate the structure and mechanism of activation we overexpressed and purified NucC in *Escherichia coli*, and then determined the crystal structure to a resolution of 1.83 Å (**Supplementary Figure S1A** and **Supplementary Table S1**). NucC is a hexamer consisting of two pairs of tightly packed trimers that pair face-to-face and bring pairs of active sites from opposite trimers together in close proximity (**Figure 1C-D**).

**Figure 1.**
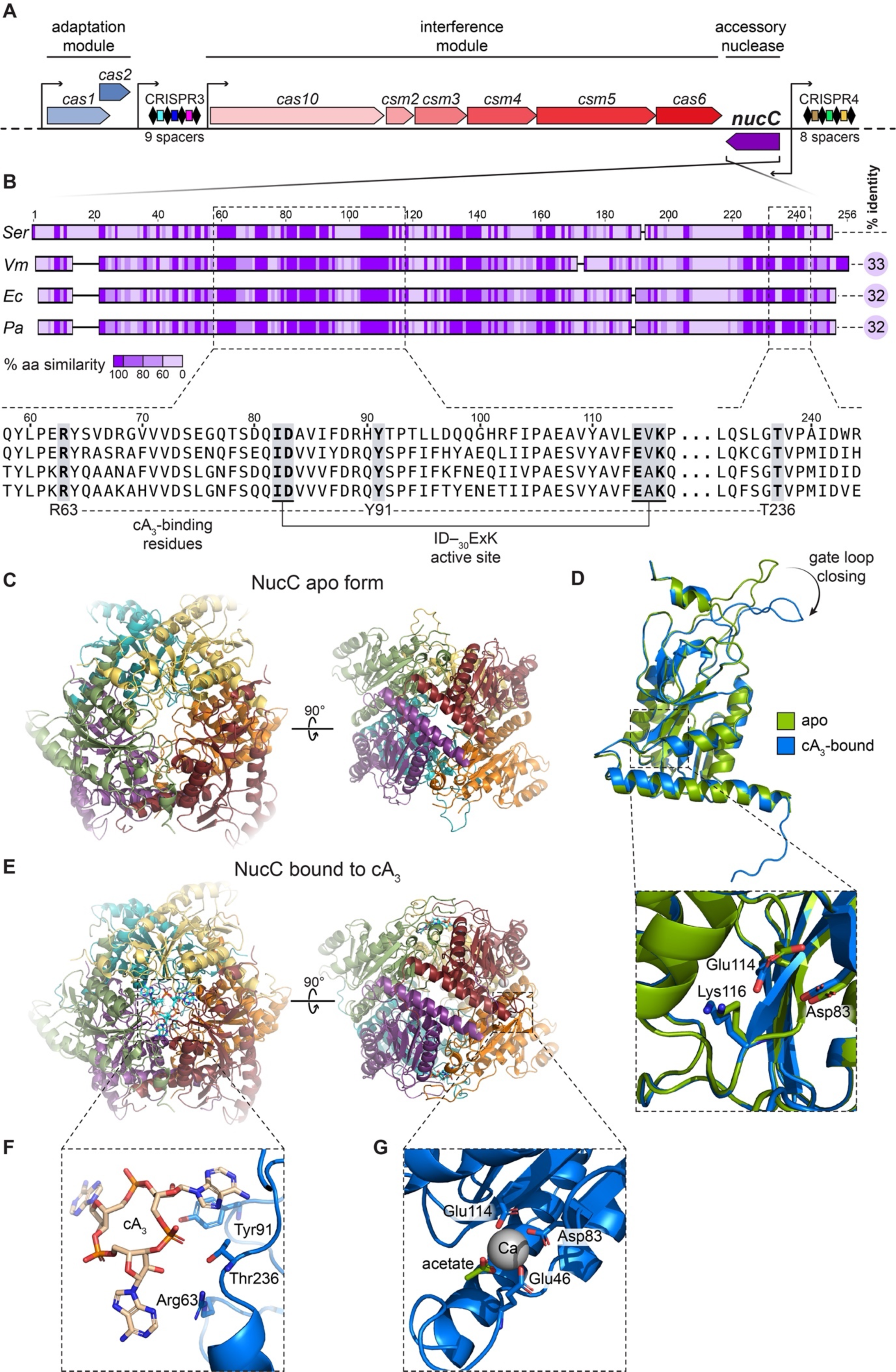
*Serratia* NucC forms a hexamer that binds cA3. (**A**) Schematic of the type III-A CRISPR– Cas operon from *Serratia* sp. ATCC 39006. Coloured boxes = spacers, black diamonds = repeats, black arrows = promoters. (**B**) Multiple Sequence Alignment (MUSCLE) of NucC homologs: *Ser*, *Serratia* type III-A CRISPR–Cas-associated NucC; *Vm*, *Vibrio metoecus* sp. RC341 type III-B CRISPR–Cas-associated NucC; *Ec*, *E. coli* MS115-1 CBASS-associated NucC; *Pa*, *P. aeruginosa* ATCC27853 CBASS-associated NucC. Numbering refers to the amino acid position in *Serratia* NucC. The percentage of amino acid similarity is indicated in different shades of purple (Score Matrix: Blosum62; Threshold: 1). The percentage of protein sequence identity is indicated for each protein sequence compared to *Serratia* NucC. Conserved cA3-binding residues (R63, Y91 and T236) and the active site motif (ID–30ExK, where ‘x’ represents any amino acid) are highlighted in grey. (**C**) Crystal structure of the *Serratia* NucC apo form at 1.83 Å. Individual monomers are coloured differently. (**D**) Conserved nuclease active site of NucC. Conformational transitions following superposition of apo (green) and cA3- bound (blue) states of the monomeric NucC monomer. (**E**) Crystal structure of *Serratia* NucC bound to cA3 at 1.48 Å. Individual monomers are differently coloured and cA3 is orange and cyan. (**F**) Interactions of conserved residues R63, Y91 and T236 with cA3. (**G**) Interactions of residues E46, D83 and E114 with Ca^2+^. Crystallographic information is reported in **Supplementary Table S1**.

Many type III accessory nucleases encode proteins containing an N-terminal CARF (CRISPR– Cas Associated Rossmann Fold) domain that binds cOA messengers and activates a variety of C-terminal effector domains (Makarova et al., 2014; Makarova et al., 2020a). In contrast, NucC proteins do not have a CARF domain despite also binding cOAs (Lau *et al*., 2020). To investigate cOA binding, we solved the co-crystal structure of NucC in the presence of its putative ligand (cA3) to a resolution of 1.48 Å (**Figure 1E** and **Supplementary Figure S1B**). NucC forms a symmetry related hexamer with two cA3 molecules. The structure revealed that a single cA3 molecule binds in a 3-fold symmetric allosteric pocket in the centre of each NucC trimer and interacts with the conserved cA3-binding residues R63, Y91 and T236 (**Figure 1B,F**). Comparison of the apo and cA3-bound structures revealed that cA3 binding induces a large conformational change in a hairpin loop (**Figure 1D**), and the closure of all three loops over the pocket completely encloses the bound cA3 (**Figure 1E**). In summary, NucC forms a hexameric structure that we predict, becomes activated upon binding cA3 and cleaves dsDNA.

### *Serratia* NucC is activated by cA3 and degrades double-stranded DNA *in vitro*

NucC homologues were previously shown to cleave plasmid and synthetic DNA *in vitro* when activated by cA3 (Grüschow *et al*., 2021; Lau *et al*., 2020). Given the predicted nuclease activity of *Serratia* NucC and its role in jumbo phage immunity (Malone *et al*., 2020), we tested its ability to degrade different nucleic acids *in vitro* in the presence of cA3. NucC degraded dsDNA when incubated with cA3 and was Mg^2+^-dependent (**Figure 2A-D**). In our crystal structures we observe a metal in the active site coordinated by the conserved acidic residues E46, D83 and E114, which we have proposed is a Ca^2+^ ion from the crystallization solution (**Figure 1G**). Notably, NucC degraded both purified *Serratia* and jumbo phage (PCH45) genomic DNA (gDNA) upon activation by cA3, resulting in a smear of smaller DNA products on the gels (**Figure 2A-B**). DNA degradation by NucC was dependent on NucC and cA3 concentration, and was observable within one minute (**Supplementary Figure 2A-C**). Mutation of predicted key nuclease active site residues (D83N, E114N and K116L) abolished the DNase activity (**Figure 1B, 1E, 2A and Supplementary Figure S2D**). Moreover, NucC was active against both supercoiled plasmid DNA and a linear PCR product (**Figure 2C-D**).

**Figure 2.**
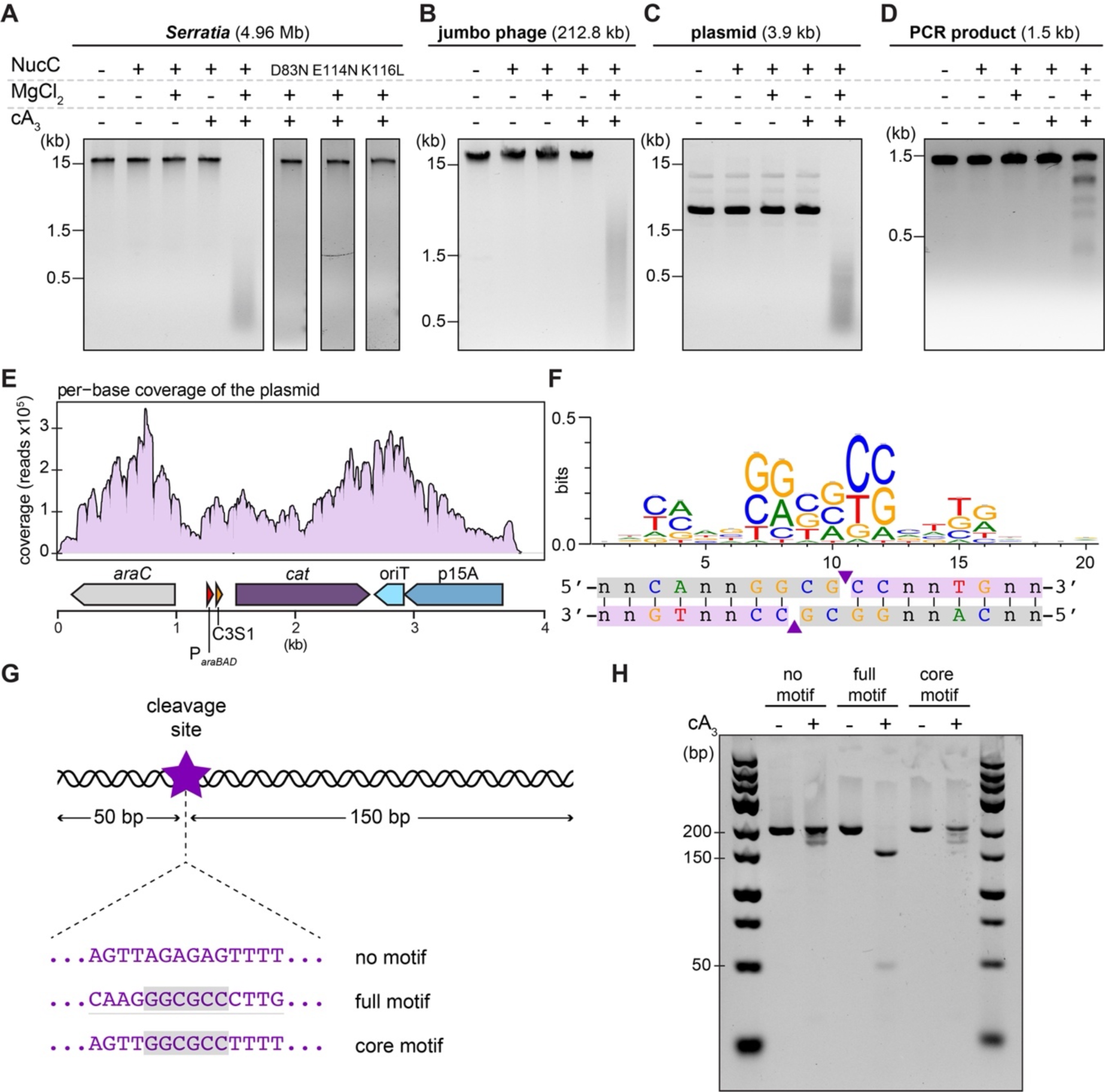
*Serratia* NucC is a cA3-activated dsDNase able to degrade *Serratia* and jumbo phage genomes *in vitro.* *In vitro* NucC cleavage of (**A**) *Serratia* and (**B**) jumbo phage (PCH45) gDNA, (**C**) plasmid and (**D**) a PCR product after 60 min incubation is dependent on MgCl2 and cA3. In (A), NucC active site mutants D83N, E114N and K116L were tested alongside the wild-type protein. (**E**) Distribution of *in vitro* NucC cleavage sites in the plasmid pPF1043 based on deep-sequencing of 5′- ends of DNA degradation products (n = 3,807,021 cleavage sites). (**F**) Cleavage site preference of NucC (WebLogo) from *in vitro* degradation products from (E), where ‘n’ represents any nucleotide. Light purple indicates the sequenced strand and purple arrows indicate the cleavage position between nucleotide (nt) positions 10 and 11. Mapping, per-base coverage and WebLogo summaries are reported in **Supplementary Tables S2**, **S3** and **S4**, respectively. (**G**) Schematic of the 200-bp synthetic dsDNA products used to verify the NucC cleavage site (purple star), with or without the specified core (shade) or full sites. The 50- and 150-bp products expected upon cleavage by NucC are indicated. (**H**) *In vitro* NucC cleavage of the synthetic oligonucleotides described in (G).

The distinct PCR product degradation pattern (**Figure 2D**) suggested that NucC might have preferred cleavage site(s). To examine sequence-specificity, we incubated a plasmid with NucC (**Supplementary Figure S2E**) and deep sequenced the resulting short fragments. The 5′-end mapping of the sequencing reads showed a heterogenous distribution of DNA degradation products (**Figure 2E**). Alignment of reads based on their 5’-ends revealed a variable 14-bp palindromic cleavage site with a 6-bp core sequence (consensus: CAnnGGCGCCnnTG, **Figure 2F**), suggesting a model of staggered double-strand cleavage with two base overhangs where two NucC-active sites cooperate to cleave both DNA strands. We observed enrichment of particular nucleotides at the outermost positions of the full motif (**Supplementary Figure S2F-G**). To verify the NucC cleavage site directly, *in vitro* cleavage assays were performed with 200-bp dsDNA fragments that contained the preferred cleavage motif with either the core or full motif (**Figure 2G**). Cleavage at the predicted site would generate 50- and 150-bp fragments. NucC specifically cleaved dsDNA only when it contained the full predicted motif sequence and upon activation by cA3 (**Figure 2H**). The diversity within the motif indicates that NucC likely cleaves additional motif variants (**Figure 2F**). Together, the NucC hexamer is activated by cA3 and cleaves double-strand DNA with some sequence-specificity.

### Type III activation of NucC triggers cell death by bacterial genome destruction

We predicted that invader RNA recognition by type III CRISPR–Cas complexes produces cA3 signals that activate NucC *in vivo* and causes growth inhibition by indiscriminate bacterial genome degradation. To explore this hypothesis, we first examined the effects of type III interference on cell growth in a phage-independent manner to discriminate NucC-dependent cell death from that induced by phage infection. To activate type III CRISPR–Cas, cA3 production and NucC, we conjugated into *Serratia* a plasmid allowing inducible target RNA expression. For this target RNA, we selected a sequence anti-sense to spacer 1 in the chromosomal CRISPR3 array which can therefore be recognised by native type III-A interference complexes (**Figure 3A**). Untargeted control plasmids (-CRISPR) conjugated efficiently into the wild-type, whereas conjugation efficiency of the targeted plasmid (+CRISPR) decreased in wild-type cells upon induction compared with the repressed control (**Figure 3B**). Despite the use of a tightly-repressed *araBAD* promoter, there was reduced conjugation efficiency for the targeted plasmid in repressed conditions. Indeed, even a single target RNA transcript from leaky expression may lead to cOA generation (Athukoralage et al., 2020a) and enable activation of NucC. In contrast to the wild-type, both the untargeted and targeted plasmids conjugated efficiently into the Δ*nucC* mutant, confirming the role of NucC in interference.

**Figure 3.**
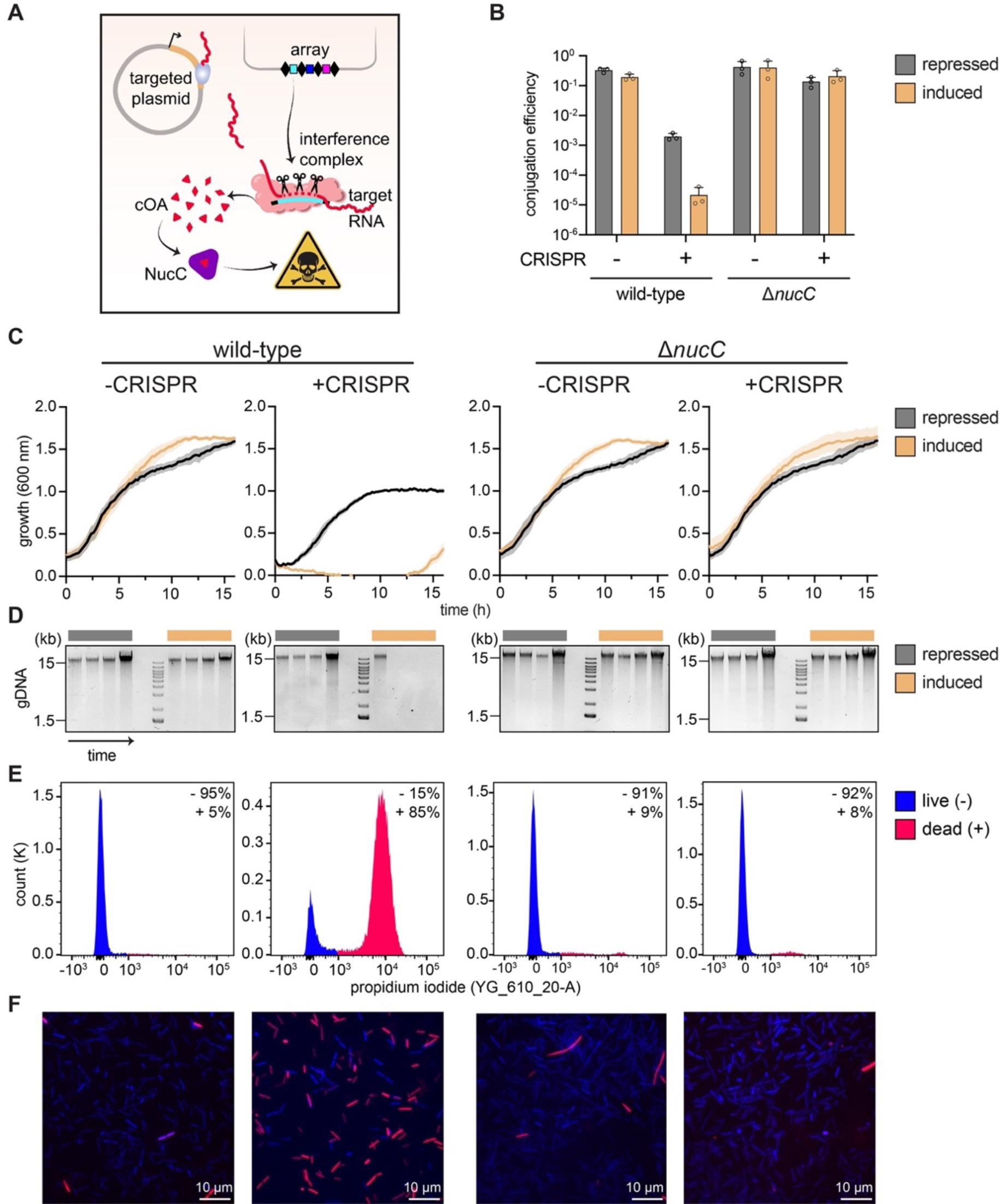
Type III CRISPR–Cas activates NucC, which degrades the bacterial genome and induces cell death. (**A**) Schematic of the plasmid targeting assay. A plasmid (+CRISPR; pPF1043) with inducible expression of an RNA targeted by the native *Serratia* type III-A system (spacer 1 of CRISPR3). RNA recognition triggers type III to synthesize cOAs that activate NucC. An empty untargeted plasmid was used as a negative control for targeting (-CRISPR; pPF781). (**B**) Conjugation efficiency assay of the targeted (+CRISPR) or untargeted (-CRISPR) plasmids into wild-type and Δ*nucC Serratia*. Cells were plated onto either glucose (glu) for RNA target repression (blue) or arabinose (ara) for RNA target induction (yellow). Wild-type and Δ*nucC Serratia* harbouring pPF781 (-CRISPR) or pPF1043 (+CRISPR) were grown under repressed (blue) or induced (yellow) conditions and monitored for (**C**) growth, (**D**) gDNA integrity (0, 0.5, 1 and 3 h post induction) and bacterial viability analysed by (**E**) flow cytometry and (**F**) confocal microscopy (1 h post induction). In (B) and (C), data represents biological triplicates as the mean ± standard deviation. In (D), (E) and (F) the data is representative of biological triplicates. The flow cytometry gating strategy and cell count data is reported in **Supplementary Figure S3B** and **Supplementary Table S5**, respectively.

Next, we assessed the effects of type III plasmid targeting on cell growth and viability. Cell growth was inhibited in wild-type cells upon induction of plasmid targeting, whereas no major effect on growth was detected with the untargeted plasmid (**Figure 3C**). Importantly, there was no inhibition of cell growth in the Δ*nucC* mutant with or without targeting, demonstrating that NucC was required for type III-dependent growth inhibition. To examine the effects of type III and NucC activation on bacterial and plasmid DNA *in vivo*, gDNA integrity and plasmid loss were examined over time after induction of plasmid targeting. Genomic DNA was degraded and plasmid DNA was purged from the wild-type host in less than 30 minutes post-induction compared with untargeted and repressed controls (**Figure 3D** and **Supplementary Figure S3A**). In contrast, genomic and plasmid DNA degradation was not detectable in the absence of NucC (**Figure 3D** and **Supplementary Figure S3A**).

Although the cell growth data demonstrated that activated type III CRISPR–Cas and NucC inhibited growth of *Serratia*, we could not discriminate between bacteriostatic (metabolic arrest) or bactericidal (cell killing) effects. To monitor bacterial viability, we assessed membrane integrity and cellular morphology from 1 to 3 h following induction of plasmid targeting. By 1 h post-induction, the type III immune response led to an increase in dead cells due to loss in membrane integrity but had little effect on cell morphology when examined by flow cytometry and confocal microscopy (**Figure 3E-F** and **Supplementary Figure S3B-E** and **Supplementary Table S5**). However, cells remained alive in the absence of type III targeting or NucC. Moreover, bacterial genome degradation upon type III and NucC activation was detected in single cells as a lower DNA staining intensity compared with controls (**Supplementary Figure S3F**). In summary, upon plasmid RNA targeting by the type III system, activated NucC degrades bacterial DNA, leading to rapid growth arrest and cell death.

### Type III jumbo phage immunity results in abortive infection

We next examined the mechanism of type III immunity and the role of NucC against the jumbo phage. Since plasmid targeting by type III activated NucC, which degraded chromosomal DNA and triggered cell death, we predicted that type III immunity against jumbo phage infection would result in abortive infection. To test this hypothesis, we used bacteria with a spacer targeting the phage capsid mRNA (**Figure 4A**) and subjected them to a single round of jumbo phage infection. Excess phage were used to ensure most bacteria were infected and cell survival quantitated by assessing whether individual infected bacteria grew into colonies. Despite providing strong immunity in efficiency of plaquing (EOP) assays (**Figure 4B**), the type III-A system with the phage-targeting spacer (+CRISPR) did not protect the viability of infected individuals, with only ∼1% surviving jumbo phage infection (**Figure 4C**). Likewise, the phage-sensitive control (-CRISPR) and Δ*nucC* mutant were dying, presumably from phage infection (**Figure 4C**). In contrast, a flagella mutant control (Δ*flhDC*) that does not produce the receptor for jumbo phage PCH45 (Smith et al., 2021) displayed complete survival, similar to the uninfected controls (**Figure 4C**). In agreement, most of the ∼1% of surviving cells detected in the phage-infected samples were spontaneous flagella mutants (**Supplementary Figure S4A**).

**Figure 4.**
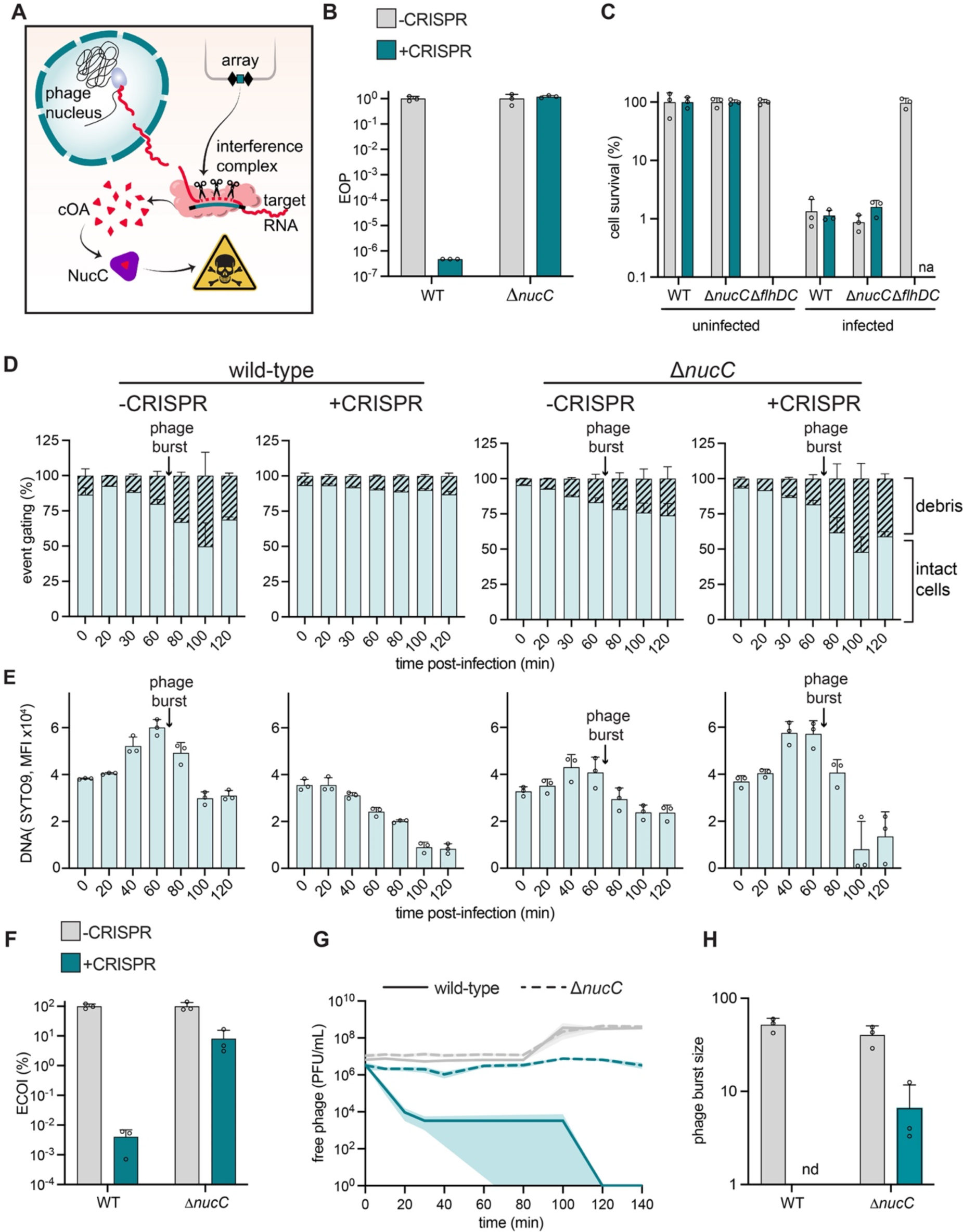
Type III immunity against jumbo phage infection causes death of the infected cell and prevents phage replication. (**A**) Schematic of the jumbo phage (PCH45) targeting assay. A type III-A crRNA is expressed from a plasmid-borne mini-CRISPR array (+CRISPR; pPF1467), which targets the jumbo phage capsid mRNA. RNA recognition activates the type III interference complex to synthesise cOAs that activate NucC. An empty vector (-CRISPR; pPF976) was used for a non-targeting negative control. (**B**) Phage resistance (efficiency of plaquing (EOP)), (**C**) cell survival, (**D**) cellular integrity, (**E**) intracellular DNA, (**F**) efficiency of centre of infection (ECOI) formation, (**G**) one-step phage infection and (**H**) average burst size for wild-type and Δ*nucC Serratia* harbouring pPF976 (-CRISPR, grey) or pPF1467 (+CRISPR, green). In (A) no individual plaques were detected in the WT +CRISPR, but lysis from without was observed. In (C) a flagella mutant (Δ*flhDC*) that lacks the jumbo phage receptor was used as a cell survival control for infection and a targeting strain was not applicable (na). Flow cytometry data for (D) and (E) are in **Supplementary Figure S4B** and **Supplementary Table S6**. No countable plaques were labelled as none detected (nd). Data in this figure represents biological triplicates plotted as the mean ± standard deviation.

Although the cell survival assay showed that cells died following infection, it could not discriminate between death due to cell suicide or phage burst. To investigate the cell death, we used flow cytometry to assess cellular integrity and DNA abundance upon jumbo phage infection and type III immunity (as in **Figure 4A**). In the absence of CRISPR immunity (- CRISPR), the number of intact cells decreased over time, reflecting cell lysis due to phage infection, whereas cells with type III immunity (+CRISPR) remained intact, indicating that cells were not lysing within 2 hours (**Figure 4D** and **Supplementary Figure S4B**). In the absence of NucC, apparent cell lysis returned, demonstrating that NucC activity prevents the jumbo phage from lysing the cell (**Figure 4D** and **Supplementary Figure S4B**). In addition, phage replication led to DNA accumulation in individual cells prior to phage burst. This accumulation in phage DNA was seen as an increase in DNA staining intensity in intact single cells of the phage-sensitive strains until the burst at around 60-80 minutes post-infection (**Figure 4E** and **Supplementary Figure S4C**). In contrast, type III immune cells exhibited NucC-dependent bacterial genome degradation (**Figure 4E** and **Supplementary Figure S4C**).

Overall, these results demonstrate that the *Serratia* type III-A system provides jumbo phage immunity via abortive infection, where infected cells die prior to completion of phage maturation and release. Therefore, we examined whether jumbo phage replication was impeded by measuring the proportion of infected cells that release phages upon type III activation (**Figure 4A**). Type III-A immunity decreased the efficiency of centre of infection (ECOI) formation in wild-type cells by >10^4^-fold (**Figure 4F**), meaning that fewer than 1 in 20,000 infected cells released at least one infectious phage. Next, we demonstrated that type III CRISPR immunity substantially reduced jumbo phage abundance in cultures throughout time when assessed in one-step growth assays (**Figure 4G** and **Supplementary Figure S4D**). Whereas the average phage burst size was around 46 in phage-sensitive cells, type III immunity completely suppressed the burst (**Figure 4H**). Importantly, complete inhibition of jumbo phage replication was NucC-dependent, as the phage could replicate in cells lacking *nucC* – albeit to a lesser extent than the phage-sensitive control (**Figure 4G-H**). This partial phage immunity in the absence of NucC may be due to contributions of Cas7 RNase or Cas10 ssDNase activity. Overall, bacterial chromosomal degradation elicited by type III immunity against jumbo phage infection results in cell death prior to completion of phage replication – the hallmark of abortive infection.

### The jumbo phage DNA-containing nucleus excludes NucC

We hypothesised that type III immunity against jumbo phage infection was provided by NucC-mediated degradation of the bacterial genome, whereas the phage DNA was protected from NucC in the nucleus-like structure. To test this, we performed phage infection assays (as in **Figure 4A**) and extracted total DNA at various times throughout a single round of phage infection. Firstly, we analysed the DNA via gel electrophoresis, which revealed no clear reduction in total DNA in phage-sensitive (-CRISPR) wild-type cells upon jumbo phage infection, indicating that the jumbo phage does not visibly degrade host DNA (**Figure 5A** and **Supplementary Figure S5A**). However, in the presence of phage targeting by the type III system (+CRISPR), high molecular weight DNA decreased, and smaller DNA degradation products became visible 40 minutes after phage addition (**Figure 5A**). In contrast, no degradation products were observed in the absence of NucC (**Figure 5A**), demonstrating that both type III CRISPR targeting and NucC were required for DNA degradation during jumbo phage infection. To determine the precise source of the degradation products (chromosomal and/or the jumbo phage DNA), we isolated the small DNA fragments for deep sequencing. At 40 min post-infection, reads mapped mainly to the *Serratia* chromosome and plasmid, but not to the jumbo phage genome (**Figure 5B** and **Supplementary Figure S5B**). At 60 min post-infection, chromosomal reads still dominated. However, there was a small increase in phage-derived reads (from ∼2% to 15% of all reads), indicating that the phage nucleus may begin to break down late in the type III-induced cell death process, leaking some phage DNA to NucC in the cytoplasm.

**Figure 5.**
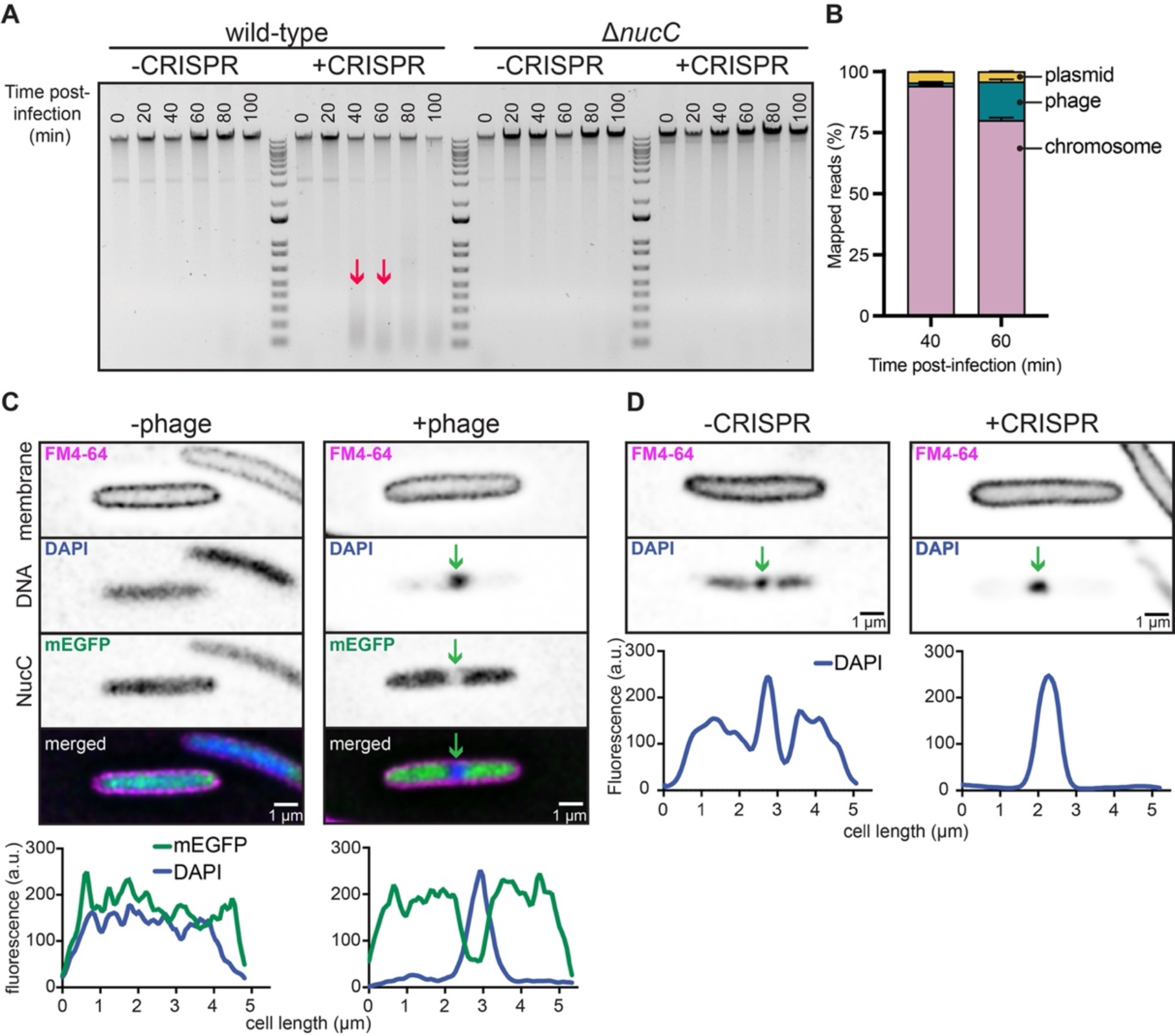
NucC cannot enter the jumbo phage nucleus but degrades the bacterial genome. (**A**) Total DNA extracted throughout a single round of jumbo phage infection and analysed via gel electrophoresis. Red arrows indicate the degradation products used for deep sequencing in (B). (**B**) Percentage of mapped reads to *Serratia* (chromosome), jumbo phage genome (phage) and plasmid DNA (pPF1467) at 40 and 60 min post-infection of wild-type +CRISPR cells resulting from deep sequencing of degradation products (red arrows) in (A). Mapping, per-base coverage and NucC motif (WebLogo) summaries are reported in **Supplementary Tables S2**, **S3, S4** and **Supplementary Figure S5E,F**. The data are biological triplicates plotted as the mean ± standard deviation. (**C**) Confocal microscopy of wild-type *Serratia* with a spacer targeting the jumbo phage (+CRISPR) in the absence (left) and presence (right) of jumbo phage infection. Membranes (magenta) and DNA (blue) were stained with FM4-64 and DAPI, respectively, while NucC tagged with mEGFP fluoresces green. Nucleus-like structures are indicated with arrows. Below, fluorescence intensity plots show the distributions of the NucC (green) and DNA (blue) across the length of single cells. (**D**) DAPI fluorescence monitored in -CRISPR and +CRISPR wild-type *Serratia* upon jumbo phage (PCH45) infection. Below, fluorescence intensity plots show the DNA distribution (blue) across the length of single cells. Images are representative of three biologically independent experiments.

Since the jumbo phage nucleus excludes the type III CRISPR–Cas complex (Malone *et al*., 2020), we hypothesized that NucC also could not access the nucleus and degrade the jumbo phage genome. To investigate this, we first generated a plasmid encoding mEGFP-tagged NucC and demonstrated that this fusion protein retained interference activity against the jumbo phage (**Supplementary Figure S5C**). Next, we studied NucC localisation by confocal microscopy during type III immunity (**Figure 4A** and **Figure 5C**). Upon phage infection, we observed circular DNA foci, consistent with phage DNA accumulation within nucleus-like structures, whereas bacterial DNA was evenly distributed in uninfected controls (**Figure 5C**). Importantly, during jumbo phage infection, NucC was localized in the cytoplasm, external to the phage DNA-containing nucleus (**Figure 5C**). By contrast, NucC was evenly distributed in the uninfected control (**Figure 5C**). We also obtained direct evidence within single cells that type III and NucC activation leads to bacterial genome degradation since bacterial DNA was undetectable upon phage targeting (+CRISPR) (**Figure 5D**). In contrast, bacterial DNA was readily detected in the cytoplasm of phage-infected cells lacking type III targeting (-CRISPR) (**Figure 5D**). In addition, jumbo phage DNA enclosed in the nucleus retained a strong fluorescent signal upon type III activation, supporting its protection from NucC activity (**Figure 5D**). Importantly, the membrane of type III immune cells (+CRISPR) remained intact compared to phage-sensitive cells (-CRISPR), indicating that cells were not lysing (**Supplementary Figure S5D**). In summary, the viral nucleus blocks NucC from accessing the jumbo phage DNA, but NucC has access to and degrades the bacterial genome, triggering abortive infection and arresting phage replication.

### Type III CRISPR provides population protection at low jumbo phage doses

We hypothesized that, despite the death of individual bacteria, type III immunity provides population-level resistance against jumbo phage infection by suppressing the phage epidemic. The inability of phage-infected cells to survive led to the prediction that population protection would fail when phages outnumber bacteria (i.e. a multiplicity of infection (MOI) > 1). To examine this hypothesis, we compared population growth of *Serratia* with a spacer targeting the jumbo phage (as in **Figure 4A**) under increasing phage pressures. Without phage targeting (-CRISPR), wild-type cells were susceptible at all phage doses, with growth inhibition faster with increased phage load (**Figure 6A**). The phages replicated extensively, reaching ∼10^10^ plaque forming units (PFU)/mL irrespective of the initial phage dose (**Figure 6A** and **Supplementary Figure S6B**). In contrast, type III immunity (+CRISPR) enabled population growth at low phage doses (MOI<1) by suppressing the phage epidemic, but when phages outnumbered bacteria (MOI>1), population growth was inhibited and phages were still detectable (**Figure 6B**). Type III immunity was lost in the absence of NucC (**Figure 6C-D**), confirming that NucC is essential for population protection and suppression of the jumbo phage epidemic. Importantly, type III immunity more rapidly inhibited culture growth upon phage infection compared with untargeted controls (**Supplementary Figure S6A**), supporting that NucC quickly induces cell death prior to phage maturation and release. In summary, type III-A immunity enables population growth under low viral load by reducing jumbo phage burden.

**Figure 6.**
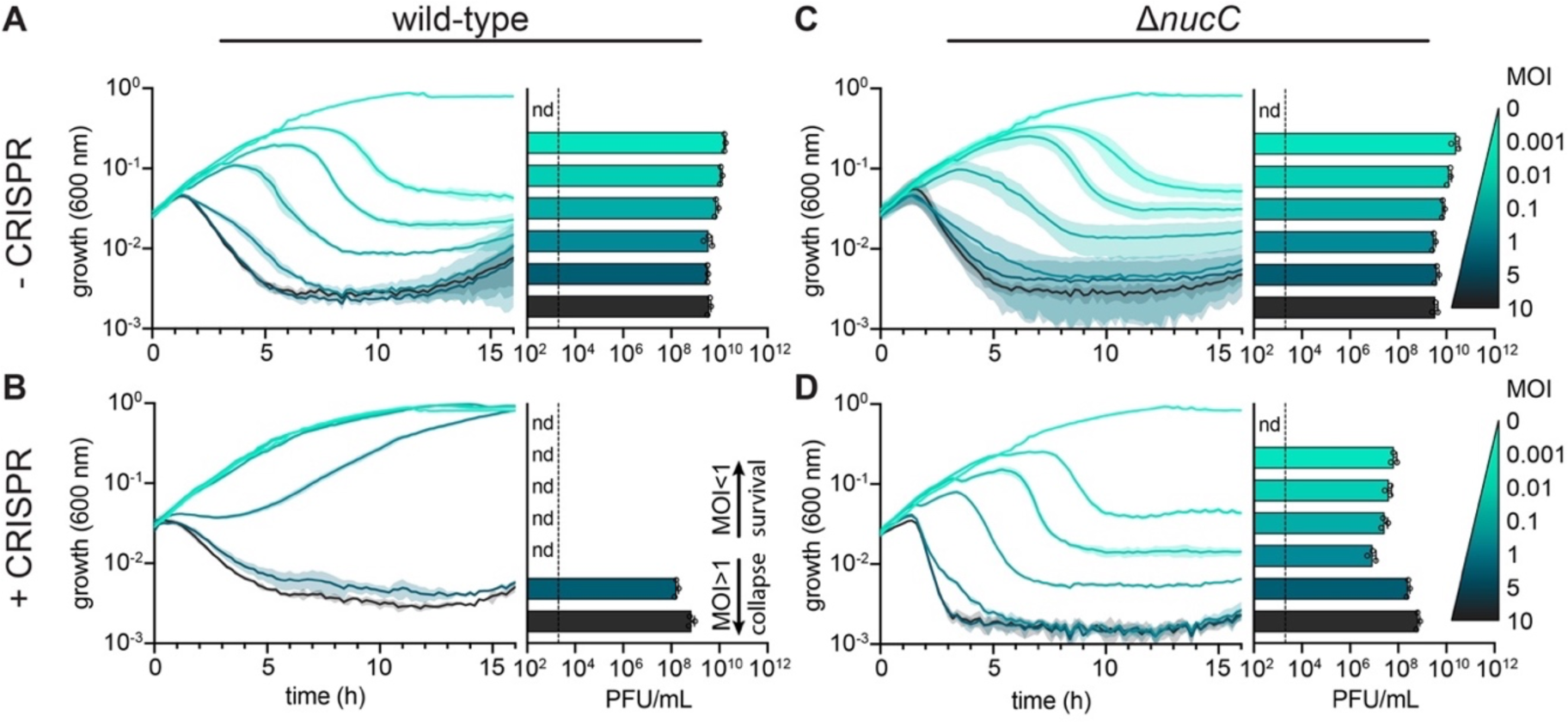
Type III CRISPR–Cas provides jumbo phage immunity by protecting the population at low phage doses and requires NucC. Infection time courses and final (16 h post-infection) phage titres were performed for wild-type and Δ*nucC Serratia* harbouring pPF976 (-CRISPR) or pPF1467 (+CRISPR), as in Figure 4A. Cultures were infected with the jumbo phage at MOIs of 0, 0.001, 0.01, 0.1, 1, 5 and 10 (coloured gradient). Final (16 h post-infection) phage titres were determined by plaque assay. Data represents biological triplicates plotted as the mean ± standard deviation. Phage infection resulting in no countable plaques were labelled none detected (nd). The limit of detection of phage titres is indicated with a dashed line.

### Type III jumbo phage immunity is widespread and can be elicited by accessory DNases and RNases

We previously demonstrated that spacers targeting nucleus-forming jumbo phages found in nature are enriched in type III compared to DNA-targeting type I CRISPR–Cas systems (Malone *et al*., 2020). Currently, the *Serratia* type III-A system with NucC is the only experimentally verified example of type III immunity against nucleus-forming jumbo phages. Therefore, we asked if NucC was specifically selected throughout evolution to provide jumbo phage immunity by type III CRISPR–Cas or whether alternative accessory nucleases could accomplish resistance. We first analysed the frequency of type III spacers targeting nucleus-forming jumbo phages associated with NucC homologs, or alternatively, with other accessory nucleases. We found that type III CRISPR–Cas systems that target nucleus-forming jumbo phages are not exclusively linked to NucC, but also to other families of accessory DNases as well as RNases such as HEPN, RelE and PD–(D/E)xK motif-containing nucleases (**Figure 7A** and **Supplementary Data File 1**).

**Figure 7.**
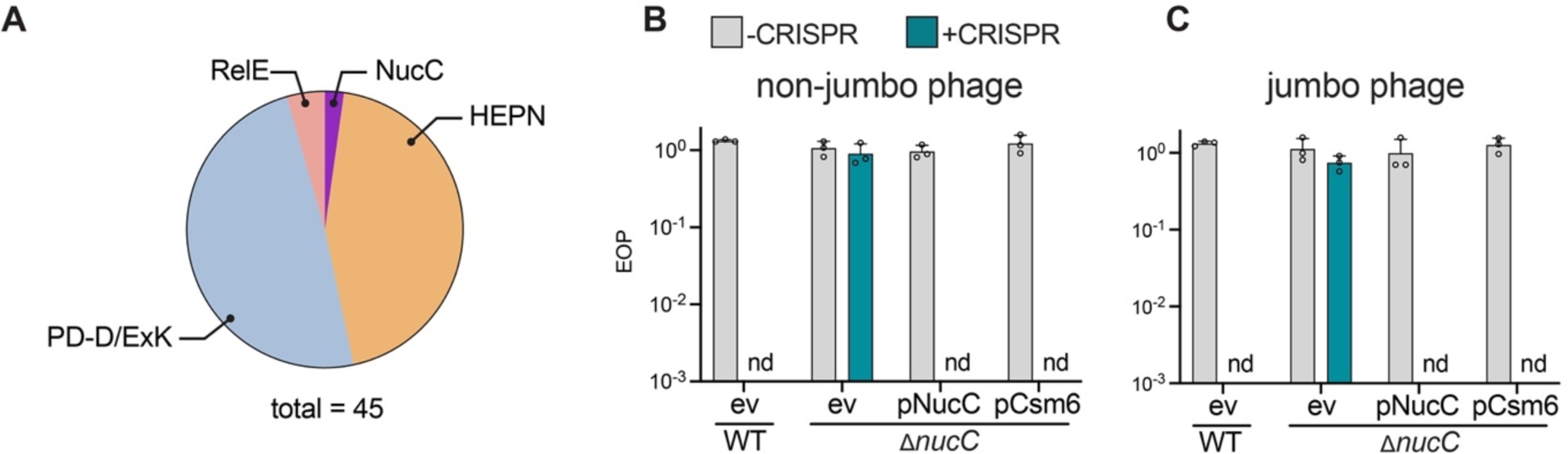
Type III jumbo phage immunity is widespread and can be elicited by accessory DNases and RNases. (**A**) Bioinformatic search for the frequency of type III spacers targeting nucleus-forming jumbo phages associating with different families of accessory nucleases. A summary of accessory nucleases identified in hosts possessing CRISPR spacers targeting nucleus-forming jumbo phages is reported in **Supplementary Data File 1**. EOP assay of (**B**) non-jumbo phage JS26 and (**C**) jumbo phage PCH45. Serial dilutions of each phage were plated on wild-type *Serratia* and △*nucC* carrying the empty vector (ev, pPF1618), pNucC (pPF2503) or pCsm6 (pPF2505). Each strain also contained a plasmid with no spacer (-CRISPR, grey, pPF976) or a spacer targeting the capsid mRNA of JS26 or PCH45 phage (+CRISPR, green, pPF1477 or pPF1467 respectively). Data shown represents biological triplicates plotted as the mean ± standard deviation. Phage infections resulting in no countable plaques were labelled none detected (nd).

The presence of natural examples of jumbo phage-targeting type III systems associated with nucleases known to target RNA led to the hypothesis that they could also provide jumbo phage immunity. Therefore, we tested whether an accessory RNA nuclease (Csm6) could protect *Serratia* from jumbo phage infection. It has been demonstrated that different nucleases can be swapped in type III-A systems to reprogram immunity (Grüschow et al., 2019), and is likely feasible due to the spectrum of different-length cOAs that Cas10 proteins produce upon target recognition (Kazlauskiene *et al*., 2017; Niewoehner *et al*., 2017; Rouillon *et al*., 2018). We exploited the *Serratia* Δ*nucC* mutant and introduced the RNase Csm6 from *Thermus thermophilus* (TtCsm6), which is activated by cA4 (Niewoehner *et al*., 2017; Niewoehner and Jinek, 2016). Given that Csm6 RNases are known to provide protection against non-jumbo phage infection, we first used a *Serratia* non-jumbo phage, the siphovirus JS26 (Jackson et al., 2019; Malone et al., 2022), as a positive control for RNase-mediated immunity. TtCsm6 and NucC both provided full immunity against JS26 in EOP assays (**Figure 7B**), which demonstrated that TtCsm6 was functional in *Serratia* and suggested that its cOA activator (cA4) was being produced by the type III-A system. Next, we examined the ability of the accessory RNase to provide jumbo phage immunity. Strikingly, TtCsm6 enabled full resistance against jumbo phage infection when compared to the phage-sensitive and empty vector controls (**Figure 7C**). In conclusion, these findings indicate that, in addition to DNases such as NucC, type III accessory RNases, such as Csm6, can provide jumbo phage immunity.

## DISCUSSION

Jumbo phages have large genomes and many undergo a complex development pathway within their bacterial hosts that includes the formation of a nucleus-like structure where DNA replication and transcription take place (Chaikeeratisak *et al*., 2017a; Chaikeeratisak *et al*., 2017b; Malone *et al*., 2020). These structures provide an anti-defence role by physically protecting the phage genome from DNA targeting defences including CRISPR–Cas (types I, II and V) and restriction–modification systems (Malone *et al*., 2020; Mendoza *et al*., 2020). In contrast, type III CRISPR–Cas systems provide bacterial immunity against jumbo phage infection and there is bioinformatic evidence the phage–host arms race has selected for spacers against jumbo phages in type III arrays (Malone *et al*., 2020). Despite these recent findings, the mechanism of type III immunity against jumbo phages remained unclear. Here we have defined how a type III CRISPR–Cas system provides protection against a nucleus-forming jumbo phage (**Figure 8**).

**Figure 8.**
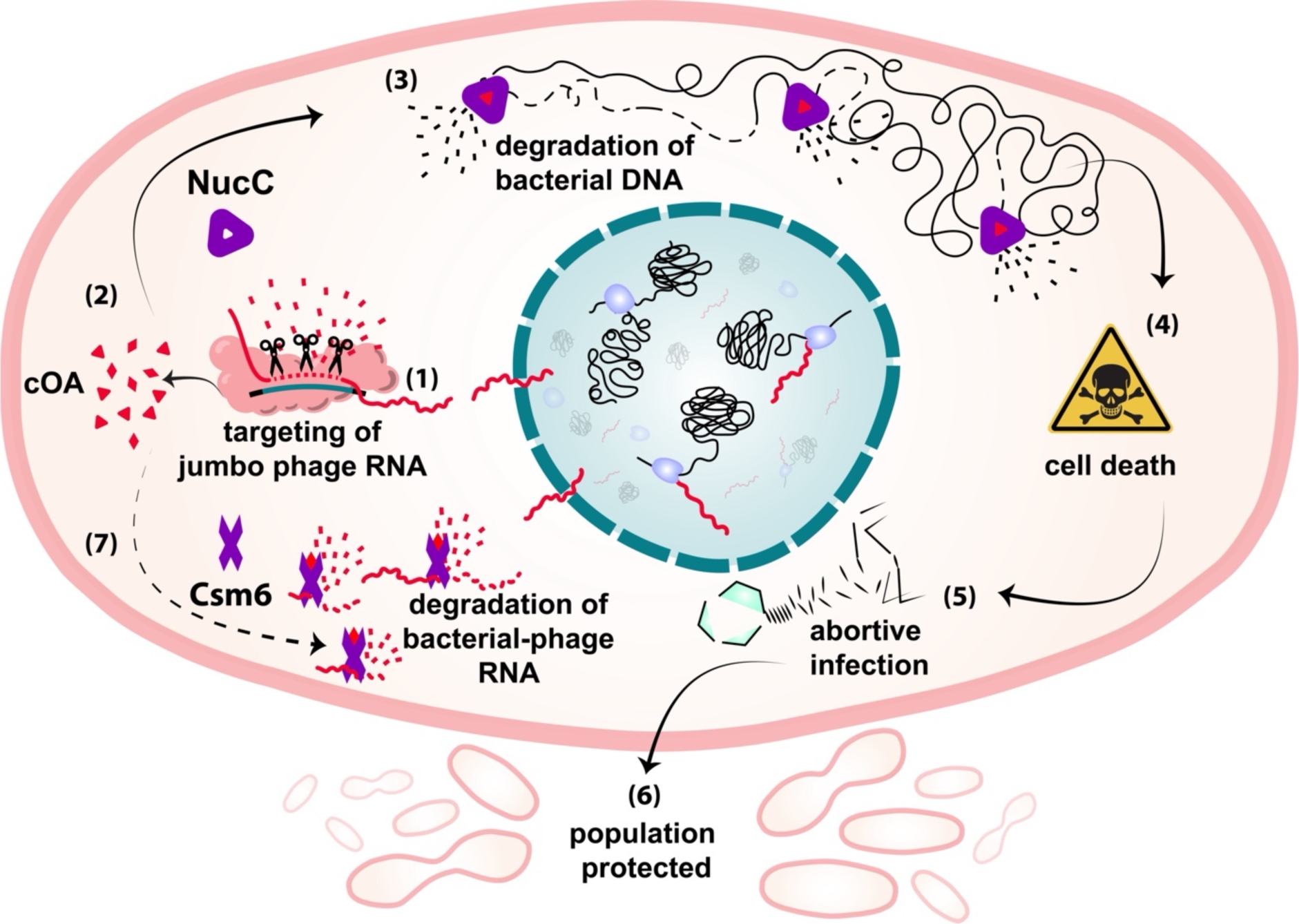
Model of type III jumbo phage immunity. Phage mRNA exiting the jumbo phage nucleus is recognised by the type III Cas complex (1), which produces cA3 (2) that activate NucC to degrade host DNA (3), eliciting cell death (4). Consequently, phage maturation and release are suppressed (5) and the clonal bacterial population is protected (6). Alternatively, type III jumbo phage immunity can be also elicited by accessory RNases such as Csm6 (7).

We demonstrate that the type III CRISPR–Cas response against jumbo phages occurs via abortive infection (Abi) in the following manner in *Serratia*. Jumbo phage (PCH45) infection is recognised by the type III complex with crRNAs matching viral mRNAs that exit the phage nucleus for translation (Malone *et al*., 2020) (**Figure 8**; step 1). Recognition of phage mRNAs results in cOA production (including cA3 and cA4) by the palm domain of Cas10 (**Figure 8**; step 2). The cA3 secondary signals are bound by hexameric NucC and leads to activation of this nuclease (**Figure 8**; step 3). Like the type III complex itself, NucC is unable to access the phage nucleus and is confined to the bacterial cytoplasm. Although excluded from the phage DNA, NucC rapidly degrades the bacterial chromosome, which results in bacterial death (**Figure 8**; steps 3 & 4), but the cells remain intact longer than if viral infection proceeded uninterrupted. The inhibitory effect of NucC blocks phage production due to the death of infected cells prior to completion of phage replication and maturation (**Figure 8**; step 5). At low viral loads, this Abi response protects the bacterial population from the jumbo phage by suppressing the phage epidemic and reducing viral spread to neighbouring cells (**Figure 8**; step 6). We also show that type III systems with spacers matching jumbo phages can harbour alternative accessory nucleases and that an RNase (Csm6) can function in place of NucC to elicit immunity against a nucleus forming phage (**Figure 8**; step 7).

Abortive infection is a phage resistance strategy elicited by a diverse range of defence systems (Isaev et al., 2021; Lopatina *et al*., 2020; Zeng et al., 2022). Recently, multiple CRISPR–Cas systems were shown to also act through Abi. For example, the type III accessory ribonucleases Csm6 and Csx1 (Foster et al., 2020; Jiang *et al*., 2016; Røstol and Marraffini, 2019) as well as the *Listeria seeligeri* type VI effector protein Cas13a (Meeske et al., 2019) indiscriminately degrade RNA, leading to cell dormancy and sometimes death. In addition, the *Pectobacterium atrosepticum* type I-F system provides population protection by aborting virulent phage infection (Watson et al., 2019). Abi generally occurs when the first lines of defence cannot inactivate the phage and the phage has proceeded into later stages of its replication cycle (Dy et al., 2014; Lopatina *et al*., 2020). Since DNA-targeting CRISPR– Cas systems are circumvented by nucleus-forming jumbo phages (Malone *et al*., 2020; Mendoza *et al*., 2020), type III immunity can function as a last resort to reduce the spread of the viral epidemic through the bacterial population. We demonstrate the importance of NucC in triggering cell death due to degradation of the bacterial chromosome since the phage DNA is inaccessible. This cell death response bears similarity to the consequences of CBASS activation of the Cap4 nuclease (Lowey et al., 2020). In contrast, in response to non-jumbo phages, type III accessory RNases can reversibly arrest growth. Reversal from dormancy occurs once phage DNA is cleared by Cas10, which also suppresses new viral mRNA, leading to a loss of new cOA production. Existing cOAs are cleaved by ring nucleases and the overall loss of secondary messengers results in accessory RNase deactivation (Athukoralage and White, 2022).

*Serratia* NucC is activated by binding cA3 and cleaves dsDNA efficiently with a degree of sequence-specificity and we have defined these sites *in vitro* and *in vivo*. In contrast, most type III accessory nucleases cut randomly, except for Card1 (Røstol *et al*., 2021). Unlike other NucC homologs (Lau *et al*., 2020), *Serratia* NucC is not activated through ligand-dependent oligomerization and exists as hexamers in apo and cA3-bound states. The conformational changes that lead to NucC activation are unclear, but we predict that cA3 binding positions or stabilizes its active sites for catalysis. The Cas10 palm domain of the *Serratia* type III-A complex synthesises at least cA3 and cA4 since phage resistance was provided when coupled with NucC (cA3-activated) or TtCsm6 (cA4-activated). This emphasises the flexibility provided by type III systems due to production of a cOA spectrum upon target recognition which enables the interchangeability of accessory effectors responsive to different cOAs (Grüschow *et al*., 2019; Shah et al., 2019). The ability of type III systems to readily swap accessory effectors is likely driven in part by the pressure imposed by phage anti-CRISPR ring nucleases, which degrade specific cOAs (Athukoralage et al., 2020b).

In conclusion, we have uncovered how type III CRISPR–Cas, in combination with the NucC accessory nuclease, provides protection against nucleus-forming jumbo phage infection. The inaccessibility of jumbo phage DNA and the Abi response of type III immunity raise the interesting question of how spacers targeting jumbo phages are acquired and whether this occurs from RNA or DNA. Nucleus-forming jumbo phages can evade a wide variety of DNA-targeting defence systems, and this quality makes them excellent candidates for phage-based therapies against bacterial infections in clinical settings. However, their vulnerability to type III immunity means we must find natural – or develop engineered – phages capable of evading RNA-targeting systems to increase the therapeutic potential of jumbo phages.

## MATERIAL AND METHODS

### Bacterial strains and growth conditions

Bacterial strains and phages used in this study are summarised in **Supplementary Table S7**. Unless otherwise noted, *Escherichia coli* and *Serratia* sp. ATCC 39006 strains were grown at 37°C and 30°C, respectively, either in lysogeny broth (LB) at 180 rpm or on LB-agar (LBA) plates containing 1.5% (w/v) agar. Minimal media contained 40 mM K2HPO4, 14.6 mM KH2PO4, 0.4 mM MgSO4, 7.6 mM (NH4)2SO4 and 0.2% (w/v) or 2% (w/v) ᴅ-glucose (glu). Phage buffer contained 10 mM Tris-HCl (pH 7.4), 10 mM MgSO4 and 0.01% (w/v) gelatine. When applicable, antibiotics and supplements were added at the following concentrations: ampicillin (Ap), 100 µg/mL; chloramphenicol (Cm), 25 µg/mL; kanamycin (Km), 50 µg/mL; gentamicin (Gm), 15 mg/mL; tetracycline (Tc), 10 µg/mL; δ-aminolevulinic acid (ALA), 50 µg/mL; isopropyl β-D-1-thiogalactopyranoside (IPTG), 50 µM; glu, 0.5% (w/v); L-arabinose (ara), 0.1% (w/v). Bacterial growth was measured as the optical density at 600 nm (OD600) using a Jenway 6300 Spectrophotometer.

### DNA isolation and manipulation

Oligonucleotides used in this study are listed in **Supplementary Table S8**. Plasmid DNA was extracted from overnight cultures using the Zyppy Plasmid Miniprep Kit (Zymo Research) and confirmed by DNA sequencing. Plasmids and their construction details are listed in **Supplementary Table S9**. Restriction digests, ligations and *E. coli* transformations were performed using standard techniques. DNA from PCRs and agarose gels was purified using the Illustra GFX PCR DNA and Gel Band Purification Kit (GE Healthcare). Polymerases, restriction enzymes and T4 ligase were obtained from New England Biolabs or Thermo Fisher Scientific.

### Multiple Sequence Alignment of NucC homologs

Multiple Sequence Alignment (MUSCLE) were performed with the NucC protein sequences from *Serratia* type III-A CRISPR–Cas system, *Vibrio metoecus* sp. RC341 type III-B CRISPR–Cas, *E. coli* MS115-1 CBASS system and *P. aeruginosa* ATCC27853 CBASS system using Geneious Prime® 2022.1.1, with a Score Matrix of Blosum62 and Threshold of 1.

### Protein expression and purification

For expression of NucC, pPF2513 was transformed into *Escherichia coli* BL21 Star (DE3) and cells were grown in LB + Km to an OD of 0.6. Expression was induced with 0.5 mM IPTG and proteins were expressed for 16 h at 18°C. Cells were harvested and resuspended in lysis buffer (20 mM HEPES-NaOH pH 7.5 and 250 mM KCl). Cells were lysed by ultrasonication and the lysate was clarified by centrifugation at 20,000 ×g for 20 min. The cleared lysate was applied to a 5 mL Ni-NTA cartridge (Qiagen). The column was washed with 3 column volumes of lysis buffer and proteins were eluted stepwise with lysis buffer supplemented with 50 and 250 mM of imidazole. The fractions eluted with 250 mM Imidazole were pooled and diluted to a final concentration of 50 mM Imidazole. TEV was added in a 1:50 ratio to allow tag cleavage at 4°C overnight. The cleavage products were passed through a 5 mL Ni-NTA cartridge (Qiagen) and the column was washed with 5 column volumes of lysis buffer supplemented with 50 mM imidazole to remove the cleaved tag and the TEV protease. The flow-through and the wash fractions were pooled, concentrated and loaded onto a HiLoad 16/600 Superdex 200 (GE Healthcare) column equilibrated in size exclusion (SEC) buffer (20 mM HEPES-NaOH pH 7.5 and 250 mM KCl). Purified NucC was concentrated using a 10,000 kDa molecular weight cut-off centrifugal filters (Merk Millipore) to a final concentration of approximately 120 mg/mL and flash-frozen in liquid nitrogen. NucC for nuclease assays was purified as above by with a GE/Cytiva Ni-NTA column and with the addition of 5% (v/v) glycerol and 1 mM dithiothreitol (DTT) to the lysis and SEC buffers.

### Protein crystallisation and structure solution

Crystals of NucC were grown at 20°C using the hanging drop vapor diffusion method. NucC was mixed 1:1 with reservoir solution. The drop size was 1 µL of protein (+ligand) and 1 µL of reservoir solution, and the reservoir had 500 µL. Apo crystals were obtained using NucC at 56 mg/mL with a reservoir containing 0.1 M sodium acetate pH 5.5, 10% PEG 8000, 10% PEG 1000 and 0.8 M sodium formate. Crystals of NucC bound to cA3 (Biolog Life Science Institute GmbH & Co. KG) were obtained by co-crystallization. NucC (24.5 mg/mL) with a 1.2 molar excess of cA3 (per trimer of NucC) were mixed 1:1 with reservoir solution containing 0.1 M Tris-HCl pH 7.5, 0.2 M calcium acetate and 25% PEG MME 2000. Crystals were flash cooled in liquid nitrogen and data were collected at the beamLine PXIII of the Swiss Light Source (Paul Scherrer Institute, Villigen, Switzerland) and processed with autoproc (Vonrhein et al., 2011). Phases were obtained by molecular replacement in Phaser (McCoy et al., 2007) implemented in Phenix, using a model obtained with Phyre2 (Kelley et al., 2015) based on PDB 6Q1H. The resulting models were completed by Autobuild (Terwilliger et al., 2008) and iterative building in Coot (Emsley et al., 2010) and refined with Phenix.refine (Afonine et al., 2012). Crystallographic information is reported in **Supplementary Table S1**.

### *In vitro* Nuclease assay

Unless otherwise noted, nucleic acids (∼100 ng) were incubated with 100 nM NucC, 200 nM cA3 and 10 mM MgCl2, in 10 mM HEPES-NaOH pH 7.5, 100 mM KCl, 5% (v/v) glycerol and 1 mM DTT. The total reaction volumes were 8 µL and were incubated at 30°C for 30 min. The samples were loaded on a 1.2% agarose gel made up in 1× TAE Buffer (Tris-acetate-EDTA), run for 40 min at 120 V and stained with ethidium bromide (EtBr).

### Isolation of *in vitro* degraded plasmid DNA

DNA was degraded by NucC *in vitro* as described above. Degraded fragments were purified with a Pippin Prep (Sage Science) using Range Mode to isolate DNA (100-400 bp) from a 2% agarose gel with EtBr staining. DNA eluted from the Pippin Prep was further cleaned and concentrated using SPRIselect left-sided size selection (2x concentration). DNA was eluted into 18 µL TE buffer and quantified using the Qubit dsDNA HS Assay kit (Thermo Fisher Scientific). This *in vitro* degraded DNA was then used to generate DNA sequencing libraries.

### DNA library preparation and sequencing

DNA sequencing libraries were prepared using the Accel-NGS 1S Plus DNA Library Kit (Swift Biosciences) according to the manufacturer’s instructions. Because samples were degraded either *in vivo* (phage infection samples) or *in vitro* (pPF1043 plasmid degradation), no DNA shearing was performed. The input DNA for each library was between 20-50 ng, and 8 cycles of indexing PCR was performed using the Accel-NGS 1s Unique Dual Indexing Kit. Final libraries were eluted in TE Buffer (Low EDTA – Swift Biosciences), quantified using the Qubit dsDNA HS Assay kit (Thermo Fisher Scientific) and fragment size distribution was determined using a Bioanalyzer High Sensitivity DNA Chip (Agilent). Libraries were diluted to 10 nM and pooled in equal ratios. The pool was then sequenced at Otago Genomics Facility (OGF) using a MiSeq Reagent kit v3 (150 cycle) to generate 2×75 bp paired end reads. Demultiplexing based on index and fastq file generation was performed by OGF as part of the Illumina MiSeq Local Run Manager standard workflow. Approximately 27 million clusters (91.7%) passed filter with an average quality score of 36.8.

### Sequencing data QC, read mapping and coverage estimation

Fastq file quality was assessed using FastQC (Andrews, 2010). The first 15 nt of Read 2 were trimmed using cutadapt (Martin, 2011) (-u 15) to remove the low complexity tail added as part of the Accel-NGS 1S Plus DNA Library Kit workflow. Reads were also filtered (-m 61) to discard those <61 nt. FastQC was re-run on trimmed samples to ensure tail removal. Reads were then mapped to the reference genome(s) using bowtie2 (Langmead and Salzberg, 2012) default parameters, specifying paired-mate mapping (--no-mixed). For *in vivo* degradation libraries, reads were mapped to a combined reference (LacA, PCH45 and pPF1467) built using bowtie2-build. For the *in vitro* degradation sample, reads were mapped to a single reference (pPF1043) built using bowtie2-build. Mapping statistics are provided in **Supplementary Table S2**. Following mapping, SAM files were converted to BAM files using SAMtools (Li et al., 2009). Average and per-base coverage was calculated (for each reference) from indexed BAM files using mosdepth (Pedersen and Quinlan, 2018). Coverage estimates are outlined in **Supplementary Table S3**.

### NucC cleavage site preference search

To generate a list of sequences to search for NucC cleavage site preference, 20 nt surrounding the first mapped base of Read 1 (9 bases upstream and 10 bases downstream of the 5’ end) were extracted as FASTA files from BAM alignments using BEDtools (Quinlan and Hall, 2010). Only Read 1 was used in the analysis, as the 5’ end of Read 2 contains a variable length low-complexity tag (introduced by the Accel-NGS 1S Plus DNA Library Kit workflow) which required trimming. Therefore, the potential cleavage position in Read 2 is ambiguous. FASTA files were then used to search for a motif for potential NucC recognition or cleavage site preferences using WebLogo (Crooks et al., 2004), where the full set of available sequences was used. Details of sequence numbers used to build motifs is outlined in **Supplementary Table S4**.

### Conjugation efficiency assay

Conjugation efficiency was assessed in a similar manner to that described previously (Patterson et al., 2015; Patterson *et al*., 2016; Richter et al., 2014). *E. coli* ST18 was the donor for the conjugation of untargeted control (-CRISPR, pPF781) and type III-A (+CRISPR, pPF1043) targeted plasmids. Plasmid pPF1043 expresses a transcript protospacer targeted by spacer 1 from CRISPR3 (type III-A), and a P*araBAD* inducible promoter controls the expression of the targeted transcript. Recipients were wild-type and Δ*nucC* (PCF686) *Serratia* strains. Strains were grown overnight in triplicate in LB for recipients, and with the addition of Cm and ALA for donor strains. One mL of overnight culture was pelleted and washed twice with LB and ALA to remove antibiotics. Pellets were resuspended in 0.5 mL of LB with ALA and glu and the OD600 was adjusted to 1. Donors and recipients were mixed in a 10:1 ratio respectively, and 10 μL was spotted on LBA with ALA and glu and incubated at 30°C for 24 h. Next, the mating spots were scrapped with a sterile loop and resuspended in 0.5 mL PBS and dilution series were plated onto LB for recipient counts or with the addition of Cm for selection of transconjugant counts. Plates also contained either glu or ara to determine conjugation under repressed or induced expression of the targeted transcript. Conjugation efficiency was calculated as the ratio of transconjugants per recipients.

### Plasmid targeting growth curves

Triplicate cultures of wild-type and Δ*nucC* (PCF686) *Serratia* harbouring the targeted plasmid (+CRISPR) or the untargeted control pPF781 (- CRISPR) were grown overnight with Cm and glu. Cultures were then incubated in 1.2 mL LB, Cm, and glu in 96-well plates from a starting OD600 ≈ 0.05 until the cultures were in the exponential phase (OD600 ≈ 0.3). Then cultures were pelleted by centrifugation, washed twice with LB and resuspended in LB with either glu or ara, from which 200 μL were transferred to a 96-well plate and incubated in a microplate reader for 16 h at 30°C with 240 rpm shaking. OD600 was monitored every 12 min.

### Plasmid targeting sample collection for genome and plasmid integrity analyses

Triplicate cultures of wild-type and Δ*nucC* (PCF686) *Serratia* harbouring the targeted plasmid pPF1043 (+CRISPR) or the untargeted control pPF781 (-CRISPR) were grown overnight with Cm and glu. Cultures were then incubated in 25 mL of LB, Cm, and glu in 250-mL flasks from a starting OD600 ≈ 0.05 until the cultures were in the exponential phase (OD600 ≈ 0.3). Then cultures were pelleted in a microcentrifuge, washed twice with LB and resuspended in LB with either glu or ara. Cultures were incubated in the same conditions, while OD600 was measured and 2x 1 mL samples were pelleted for gDNA and plasmid extraction at 0, 0.5, 1 and 3 h post-infection. Genomic DNA was extracted from pellets using the DNeasy Blood & Tissue kit (Qiagen) per the manufacturer’s instruction. Plasmid DNA was extracted from pellets using the Zyppy Plasmid Miniprep Kit (Zymo Research). Then 5 μL of gDNA or 20 μL of plasmid were loaded onto a 1% agarose gel made up in 1× TAE buffer, run for 40 min at 100 V and stained with EtBr.

### Cell viability assay during plasmid targeting

Triplicate cultures of wild-type and Δ*nucC* (PCF686) *Serratia* harbouring the targeted plasmid pPF1043 (+CRISPR) or the untargeted control pPF781 (-CRISPR) were grown overnight. Aliquots of overnight cultures (500 μL) were used to inoculate new universals containing 5 mL of LB with Cm and glu and incubated for 3 h. Cells were collected by centrifugation and resuspended in 5 mL of LB with ara to induce target expression. After 1 h at 30°C with shaking (160 rpm), 500 μL was pelleted by centrifugation and resuspended in 500 μL of minimal media. Cells were stained with the LIVE/DEAD BacLight Bacterial Viability Kit (Invitrogen) by adding 1 μL of the SYTO9 and propidium iodide (PI) and incubating the samples in the dark at room temperature (RT) for 5 min. Samples were washed with 1 mL minimal media and resuspended in 100 μL minimal media. Finally, 1 μL of cell suspension was added to 1 mL of 1× phosphate-buffered saline (PBS) and analysed using the LSRFortessa flow cytometer (BD Biosciences). The SYTO9 was excited using a blue laser (488 nm) and detected with a 530/30 nm bandpass filter and the PI was excited using a yellow–green laser (561 nm) and detected with a 610/20 nm bandpass filter and 20,000 events were recorded per sample using the BD FACSDiva software (v.8, BD Biosciences). Subsequent analysis was performed using FlowJo software v.10 (BD Biosciences). Cells were gated on positive SYTO9 fluorescence (total bacterial cells) and the subpopulation of dead cell was determined by median PI fluorescence intensity (MFI) of the cell populations. Three biological replicates per strain were used for each experiment, and data plotted as the mean ± SD. Cell counts of cell viability assay during plasmid targeting is reported in **Supplementary Table S5**.

### Cell viability assay

To monitor cell death upon type III-A targeting, we performed confocal microscopy of wild-type and Δ*nucC Serratia* harbouring the targeted plasmid pPF1043 (+CRISPR) or the untargeted control pPF781 (-CRISPR). Overnight cultures were grown at 30°C with Cm and glu (0.1% w/v) to repress the expression of the targeted sequence. Aliquots of overnight cultures (500 μL) were used to inoculate flasks with 5 mL of LB with Cm and glu (0.1% w/v) and incubated for 3 h. Cells were collected by centrifugation and resuspended in 5 mL of LB with ara (0.5% w/v) to induce target expression. After 1 h incubation at 30°C in shaking conditions (160 rpm), 500 μL cell culture were pelleted by centrifugation and resuspended in 200 μL of minimal media. Cells were stained with 4,6-diamidino-2- phenylindole (DAPI; 4 μg/mL) and propidium iodide (PI, 66 μg/mL) and incubated the samples in the dark at room temperature (RT) for 5 min. Stained samples were then washed with minimal medium before being resuspended in 100 μL of minimal medium. To suspend samples, 15 μL aliquots of each sample were mixed with 15 μL of molten 1.2% (w/v) agarose (in minimal media) before being sealed onto microscope slides with a coverslip.

### Confocal Microscopy and Image Analysis

Images were acquired using a CFI Plan APO Lambda ×100 1.49 numerical aperture oil objective (Nikon Corporation) on the multimodal imaging platform Dragonfly v.505 (Oxford Instruments) equipped with 405, 488, 561 and 637 nm lasers built on a Nikon Ti2-E microscope body with Perfect Focus System (Nikon Corporation). Data were collected in Spinning Disk 40 μm pinhole mode on the iXon888 EMCCD camera with x2 optical magnification using the Fusion Studio v.1.4 software. Z stacks were collected with 0.1 μm increments on the z axis using an Applied Scientific Instrumentation stage with 500 μm piezo z drive. Images were visualized and cropped using Fiji software (Windows 64-bit) and further processed using the Huygens Essential Deconvolution Wizard (Scientific Volume Imaging). Final composite images and fluorescence plot data were generated using Fiji and graphed using Prism v. 9.2.0 (GraphPad).

### Bacteriophage purification and titration

Phage stocks were prepared as described elsewhere (Watson et al., 2018). Briefly, 100 μL of overnight bacterial culture was mixed with serial dilutions of phage lysate and added to 4 mL of LBA overlay (0.35% w/v), which was then poured onto LBA plates. Plates were then incubated overnight at 30°C; plates with almost confluent lysis had the top agar scraped off and pooled together into a centrifuge tube. A few drops of chloroform (NaCO3-saturated) were added before thoroughly vortexing the mixture to lyse the cells. Finally, a centrifugation step was performed (2,000 ×g for 20 min at 4°C) to separate the virions from the cell debris. The supernatant was placed in a sterile universal for storage and phage titre was determined by pipetting 20 μL drops of serial dilutions of the phage stock in phage buffer onto a LBA overlay seeded with 100 μL of *Serratia* overnight culture. Plaques were counted after incubation overnight at 30°C, with the phage titre represented as plaque-forming units (PFUs) per millilitre (PFU/mL). Phage stocks were stored at 4°C.

### Efficiency of plaquing (EOP) assay

A soft LBA overlay (0.35% w/v) containing 100 μL of bacterial culture was poured onto a LBA plate. Serial tenfold dilutions of high-titre PCH45 phage stock (approximately 10^9^ PFU/mL) were spotted (20 μL) onto the agar overlay and plates were incubated overnight at 30°C. The efficiency of plaquing (EOP) was calculated as the ratio of PFU/mL produced on tested strains to the PFU/mL on the control *Serratia* strain.

### Cell survival

Triplicate cultures of wild-type and Δ*nucC* (PCF686) *Serratia* harbouring the anti-phage spacer-containing plasmid pPF1467 (+CRISPR) or the empty vector control pPF976 (-CRISPR) were grown overnight. The overnight cultures were subcultured into 5 mL LB from a starting OD600 of 0.05. Cells were grown until early stationary phase (OD600 of ∼0.3) before aliquots of 1 mL were transferred into two Bijou tubes. One culture was infected with PCH45 at a multiplicity of infection (MOI) of 10, while the other was mock infected with phage buffer. Cultures were incubated with shaking for 20 min for phages to adsorb, and the 1 mL was transferred into microcentrifuge tubes. Cells were pelleted, washed twice in PBS to remove unadsorbed phages, serial diluted and plated before the infected cells starting lysing (∼80 min). Cell survival was calculated as (colony forming units (CFU)/mL (infected sample)/CFU/mL (uninfected sample).

### Swimming motility plate assay

Individual colonies were picked and stabbed into Tryptone Swarm Agar (TSA) media (10 g/L Bacto tryptone, 5 g/L NaCl, 3 g/L Bacto agar). Plates were incubated at 30°C overnight, and halo formation was examined to test for swimming defects indicative of flagella mutations (jumbo phage PCH45 receptor mutants).

### Cell viability assay during phage infection

Triplicate cultures of wild-type and Δ*nucC* (PCF686) *Serratia* harbouring the anti-phage spacer-containing plasmid pPF1467 (+CRISPR) or the empty vector control pPF976 (-CRISPR) were grown overnight. The following day, 100 µL of each culture was transferred into 5 mL LB media and grown until exponential phase (∼3 h; OD600 = 0.3) at 30°C with 180 rpm shaking. Prior to the addition of phage, 200 µL of culture was removed as a pre-infection control (time 0). To the remaining culture, phage PCH45 was added to MOI of 10. Cultures were then incubated at 30°C with 180 rpm shaking. At each time point (20, 40, 60, 80, 100, and 120 min-post infection), 200 µL of culture was removed to a 1.5 mL microcentrifuge tube and centrifuged at 17,000 ×g for 1 min. The supernatant was decanted and pellets resuspended in 1 mL PBS. Cells were centrifuged at 17,000 ×g for 1 min, supernatant was removed, and pellets resuspended in 50 µL PBS. To each sample, 2 µL of a stain mix (equal parts SYTO9 and Propidium Iodide from the LIVE/DEAD^™^ *Bac*Light^™^ kit) was added. Samples were vortexed and incubated at RT for 5 min protected from light. Cells were then centrifuged at 17,000 ×g for 1 min, the supernatant removed, and pellets washed with 1 mL PBS. Washed cells were centrifuged at 17,000 ×g for 1 min, the supernatant removed, and samples resuspended in PBS (between 20 µL – 50 µL, depending on pellet density). For flow cytometry analysis, 10 µL of resuspended cells were diluted in 1 mL PBS in a round-bottom falcon tube (Corning). Samples were analyzed on a LSRFortessa (BD Biosciences). Thresholds of 200 (forward-scatter) and 500 (side-scatter) were applied to localize bacterial cells. SYTO9 was detected with 530/30 filter (488 nm blue laser) at 475 V and PI was detected with 610/20 filter (561nm yellow-green laser) at 650 V. For each sample, 10,000 events were recorded. Data was analyzed using FlowJo v10.8.1 (BD Biosciences). Median channel (530/30 and 610/20) area values were used for SYTO9 and PI fluorescence levels. Uninfected samples were used to position gates which delineate intact cells (as shown in **Supplementary Figure S3A**). Data used to construct **Figures 4D-E** is shown in **Supplementary Table S6**. Bar graphs were generated in Prism and histograms were generated in FlowJo.

### Efficiency of centre of infection (ECOI)

Triplicate cultures of wild-type and Δ*nucC* (PCF686) *Serratia* harbouring the anti-phage spacer-containing plasmid pPF1467 (+CRISPR) or the empty vector control pPF976 (-CRISPR) were grown overnight. In addition, the flagella mutant (Δ*flhDC*) that lacks the PCH45 receptor was used as a control for cell survival following phage infection. The overnight cultures were used to inoculate a 5 mL culture from a starting OD600 of 0.05. Cells were grown until early stationary phase (OD600 of ∼0.3) before PCH45 was added at an MOI of 0.1, and cultures were incubated with shaking for 20 min for phages to adsorb. Aliquots of 1 mL were extracted, cells pelleted, washed twice in PBS to remove unadsorbed phages, serial diluted and plated in top LBA with wild-type before the infected cells starting lysing (∼80 min). The PFU/mL was determined for each strain and since each plaque was formed from the phages released from an individual cell, the titre represents the number of infectious centres formed. ECOI (%) was calculated as the ratio of PFU/mL produced on tested strains to PFU/mL on the control strain, multiplied by 100.

### One-step growth curve and burst size

Triplicate cultures of wild-type and Δ*nucC* (PCF686) *Serratia* harbouring the anti-phage spacer-containing plasmid pPF1467 (+CRISPR) or the empty vector control pPF976 (-CRISPR) were grown overnight. The overnight cultures were used to inoculate a 10 mL culture from a starting OD600 of 0.05. Cells were grown until early stationary phase (OD600 of ∼0.3) before PCH45 was added at an MOI of 0.1. Cultures were incubated in the same conditions and two samples of 100 µL were removed periodically (0, 10, 20, 30, 40, 60, 80, 100, 120 and 140 min) and mixed with 900 µL (untreated) or 800 µL PBS with 20 µL of chloroform (NaCO3-saturated) (treated). Untreated samples were immediately serially diluted in LB and titrated as described above to determine the number of PFU/mL. Treated samples were immediately vortexed, and only diluted and titrated after all the untreated samples. The average burst size was calculated as the PFU/mL initially added subtracted from the PFU/mL in the second round of infection, divided by the PFU/mL initially added and results were expressed as the ratio of PFU/mL per infected cell.

### Visualization of degraded DNA upon phage infection

Wild-type and τ−.*nucC* (PCF686) harbouring either a non-targeting plasmid (pPF976) or a plasmid with a PCH45 targeting spacer (pPF1467) were grown overnight in 5 mL LB + Km (50 µg/mL) and 100 µM IPTG (for spacer induction) at 30°C with shaking at 180 rpm. The following day, strains were subcultured to a starting OD600 = 0.05 in 25 mL LB + Km (50 µg/mL) and 100 µM IPTG in 125 mL flasks. Cells were grown approximately 4 h until reaching an OD600 = 0.3. One mL of each culture was removed for gDNA extraction as a pre-infection (time 0) control. Ten mL of each culture was then removed to a universal, and phage PCH45 was added at an MOI of 10. Cultures were incubated at 30°C with 180 rpm shaking and samples were taken at the following time points: 20, 40, 60, 80, and 100 min-post infections. At each time point, 1 mL of culture was removed and pelleted at 17,000 ×g for 1 min. Supernatant was removed and pellets were washed twice with 1 mL PBS. DNA was then extracted using the DNeasy Blood & Tissue kit (Qiagen) per the manufacturer’s instruction. Briefly, each pellet was resuspended in 180 µL Buffer ATL with 20 µL Proteinase K and incubated at 56°C for 30 min. Following incubation, 4 µL RNase A (10 mg/mL) was added to each tube and incubated at RT for 5 min before proceeding with DNeasy procedure. Purified DNA was eluted with 30 µL TE buffer. Sample concentration was measured using a NanoDrop spectrophotometer and then diluted to a concentration of 20 ng/µL. For each sample, 500 ng was loaded onto a 1% agarose gel made up in 1× TAE buffer, run for 40 min at 100 V and stained with EtBr.

### Isolation of gDNA degradation products during phage infection

Triplicate cultures of wild-type and Δ*nucC* (PCF686) *Serratia* harbouring the anti-phage spacer-containing plasmid pPF1467 (+CRISPR) or the empty vector control pPF976 (-CRISPR) were grown overnight. The overnight cultures were subcultured, infected, and grown as described in the previous section - Visualization of degraded DNA upon phage infection. At each time point, (pre-infection, and 20, 40, 60, 80, and 100 min-post-infection), 1.5 mL of culture was removed and pelleted at 17,000 ×g for 1 min. Cells were washed and DNA was extracted as described above. DNA was eluted in 50 µL TE buffer. To separate intact genomic fragments from degraded DNA, a right-sided size selection was performed using SPRIselect beads (Beckman Coulter) with a 20 µL elution from the first bead addition (0.6x concentration) to recover genomic fragments, and a 35 µL elution from the second bead addition (1.2x concentration) to recover degraded fragments. To remove any carryover of intact genomic DNA, degraded fragments were further purified with a Pippin Prep (Sage Science) using Range Mode to isolate DNA (100-400 bp) from a 2% agarose gel with EtBr staining. DNA eluted from the Pippin Prep was further cleaned and concentrated using SPRIselect left-sided size selection (2x concentration). DNA was eluted into 18 µL TE buffer and quantified using the Qubit dsDNA HS Assay kit (Thermo Fisher Scientific). DNA isolated from wild-type *Serratia* with CRISPR targeting (pPF1467) at 40 min (n=3) and 60 min (n=3) post PCH45 phage infection was used to generate sequencing libraries. These time points were chosen as DNA degradation became visible 40 min-post infection (**Figure 5A**).

### NucC localization microscopy

To visualize NucC localization in wild-type *Serratia* cells, an N-terminal NucC-mEGFP fusion was expressed under control of P*araBAD* (ara-inducible). Cells harbouring the NucC-mEGFP expression plasmid (pPF2290) harboured a second plasmid, containing either a protospacer matching phage PCH45 (pPF1467) or a control plasmid (pPF976). Overnight cultures grown in LB + Km + Gm were used to seed new 25 mL cultures in 125 mL flasks at starting OD600 = 0.05. Cells were grown in LB + 0.1% ara (w/v) for NucC- mEGFP induction, 100 µM IPTG for protospacer induction, and Km + Gm for plasmid maintenance. Cultures were grown at 30°C with 180 rpm shaking for ∼3.5 h until reaching exponential phase (OD600=0.3). Cultures were then split into 5 mL aliquots in glass universals. For +phage treatments, phage PCH45 (∼1×10^11^ PFU/mL) was added at an MOI of 50. To - phage treatments, an equivalent volume of phage buffer was added. Infected and non-infected cultures were grown for 50 min at 30°C with 180 rpm shaking. Following growth, 1.5 mL of each culture was removed to a 1.5 mL microcentrifuge tube and centrifuged at 17,000 ×g to pellet cells. Each pellet was washed 2× with 500 µL minimal media. Pellets were resuspended in 34 µL minimal media, and 16 µL of stain mix (4,6-diamidino-2-phenylindole (DAPI; final 4 µg/mL) and FM4-64 (final 12 µg/mL)) was added to each sample. Samples were incubated in the dark at RT for 5 min, then centrifuged for 30 s at 17,000 ×g. Supernatant was removed, pellets washed with 500 µL minimal media centrifuged for 30 s at 17,000 ×g. Supernatant was removed, and pellets resuspended in 50 µL minimal media. Samples preparation, confocal microscopy and image analysis were performed as described above.

### Phage resistance infection time courses

Triplicate cultures of wild-type and Δ*nucC* (PCF686) *Serratia* harbouring the anti-phage spacer-containing plasmid pPF1467 (+CRISPR) or the empty vector control pPF976 (-CRISPR) were grown overnight. The overnight cultures were subcultured in 1.2 mL LB media from a starting OD600 of 0.05. Cells were grown until early stationary phase (OD600 of ∼0.3) and cultures were diluted again to an OD600 of 0.05 before PCH45 was added at a range of MOIs and growth (OD600) was monitored every 12 min for 16 h. Phage titres were determined at the end of the experiment as described earlier.

### Identification of accessory nucleases in hosts possessing CRISPR spacers targeting nucleus-forming jumbo phages

First, we used CRISPRDetect 2.4 (Biswas et al., 2016) to identify CRISPR loci in the genomes of hosts in the RefSeq v201 Archaea and Bacteria database. We then searched for matches to the spacers in a set of nucleus-forming phage contigs in the IMG_VR v3 (5.1) dataset (Roux et al., 2021). The IMG_VR contigs were assigned as belonging to nucleus-forming phages if genes encoding homologs of both the shell and tubulin proteins were identified within the corresponding vOTU, with the shell and tubulin genes identified as previously described (Malone *et al*., 2020). For each host possessing spacers targeting nucleus-forming phages, we searched for genes encoding CRISPR–Cas systems and accessory nucleases using PADLOC (Payne et al., 2022; Payne et al., 2021) and classified hits according to the presence of conserved CARF and nuclease domains (Makarova *et al*., 2020a). A summary of accessory nucleases identified in hosts possessing CRISPR spacers targeting nucleus-forming jumbo phages is reported in **Supplementary Data File 1**.

### Analysis of function of accessory nucleases

For *in vivo* expression of accessory nucleases, pPF2503 (pNucC), pPF2505 (pCsm6) or pPF1618 (empty vector) were introduced into *Escherichia coli* ST18 by heat shock transformation. Subsequently, the plasmids were moved by conjugation into *Serratia* wild-type (only empty vector control) or the *nucC* mutant (PCF686) containing a plasmid-borne type III-A spacer targeting PCH45 or JS26 (+CRISPR, pPF1467 or pPF1477, respectively) and the corresponding no spacer control (-CRISPR, pPF976). EOP assays were performed as described above.

## AUTHOR CONTRIBUTIONS

D.M.-M., L.M.M., H.G.H. and P.C.F. conceived the project with input from all authors. D.M.- M., C.G.-D., L.M.M., K.R.H. and S.A.J. generated strains and plasmids. D.M.-M., L.M.S., C.G.- D., L.M.M., H.G.H., L.F.G. and P.C.F. designed experiments. C.G.-D. solved the NucC crystal structure. D.M.-M., C.G.-D. and R.D.F. purified NucC. D.M.-M. and L.M.S. performed *in vitro* cleavage assays. L.M.S. performed NucC degradation assays by deep-sequencing. D.M.-M. and L.M.M. performed plasmid targeting experiments. D.M.-M., L.M.S. and K.R.H. performed phage infection assays. L.M.S. and L.M.M. performed flow cytometry. L.M.S., L.M.M. and L.F.G. performed confocal microscopy. S.A.J. performed bioinformatics of accessory nucleases. D.M.-M., L.M.S., C.G.-D. and P.C.F. designed figures. D.M.-M. and P.C.F. wrote the manuscript with input from all authors.

## DATA AVAILABILITY

Reference sequences are available from NCBI for *Serratia* sp. ATCC 39006 LacA: GCF_002847015.1_ASM284701v1; RefSeq: NZ_CP025085.1 and *Serratia* phage PCH45: GCA_009187865.1_ASM918786v1; GenBank MN334766.1. NucC-mediated DNA degradation sequencing is BioProject accession PRJNA846505 and BioSample accession SAMN28887774 (Reviewer link: https://dataview.ncbi.nlm.nih.gov/object/PRJNA846505?reviewer=gohaf3qp7mdvrlpgdgv85pjmuo). The crystal structures of the *Serratia* NucC in its apo (PDB ID 7ZGW) and cA3-bound (PDB ID 7ZGV) forms are available.

## ACKNOWLEDGEMENTS

This work was supported by the Marsden Fund from the Royal Society of New Zealand (Te Pūtea Rangahau a Marsden, Te Apārangi). D.M.-M. was supported by a University of Otago Doctoral Scholarship. L.M.M. was supported by the Health Sciences Career Development Award of the University of Otago. We thank the staff of the Otago Micro and Nanoscale Imaging and Otago Genomics Facilities for assistance with flow cytometry, confocal microscopy and genome sequencing. We thank Vincent Olieric, Takashi Tomizaki and Meitian Wang (Swiss Light Source, Paul Scherrer Institute, Villigen, Switzerland) for assistance with crystallographic data collection and analysis. We are grateful to Martin Jinek for support, discussions and access to resources. I’ll be fine with whatever you decide.We thank members of the Fineran laboratory for helpful discussions. We are especially thankful to Nils Birkholz for valuable input.

