## Supplementary Information for "Type III CRISPR–Cas provides resistance against nucleus-forming jumbo phages via abortive infection"

### SUPPLEMENTARY MATERIAL

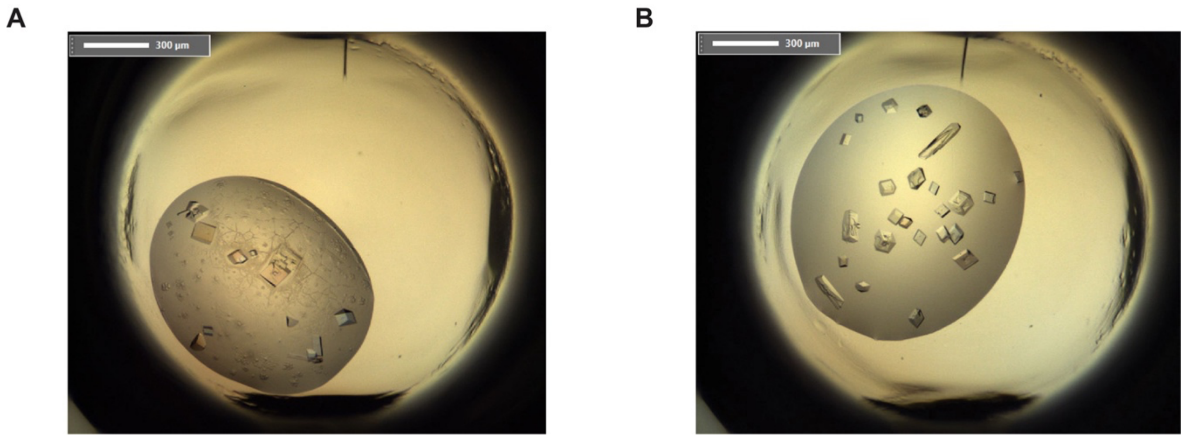


**Supplementary Figure S1. NucC crystallization.** (**A**) Crystallization of *Serratia* NucC apo form at 1.83 Å; NucC = 56 mg/mL, 0.1 M Na acetate pH 5.5, 10% PEG 8K, 10% PEG 1K, 0.8 M Na formate. (**B**) Co-crystallization of *Serratia* NucC bound to $cA_3$ at 1.48 Å; NucC = 24.5 mg/mL (+1.2x $cA_3$/trimer), 0.1 M Tris pH 7.5, 0.2 Ca acetate, 25% PEG MME 2K. Crystallographic information is reported in **Supplementary Table S1**.

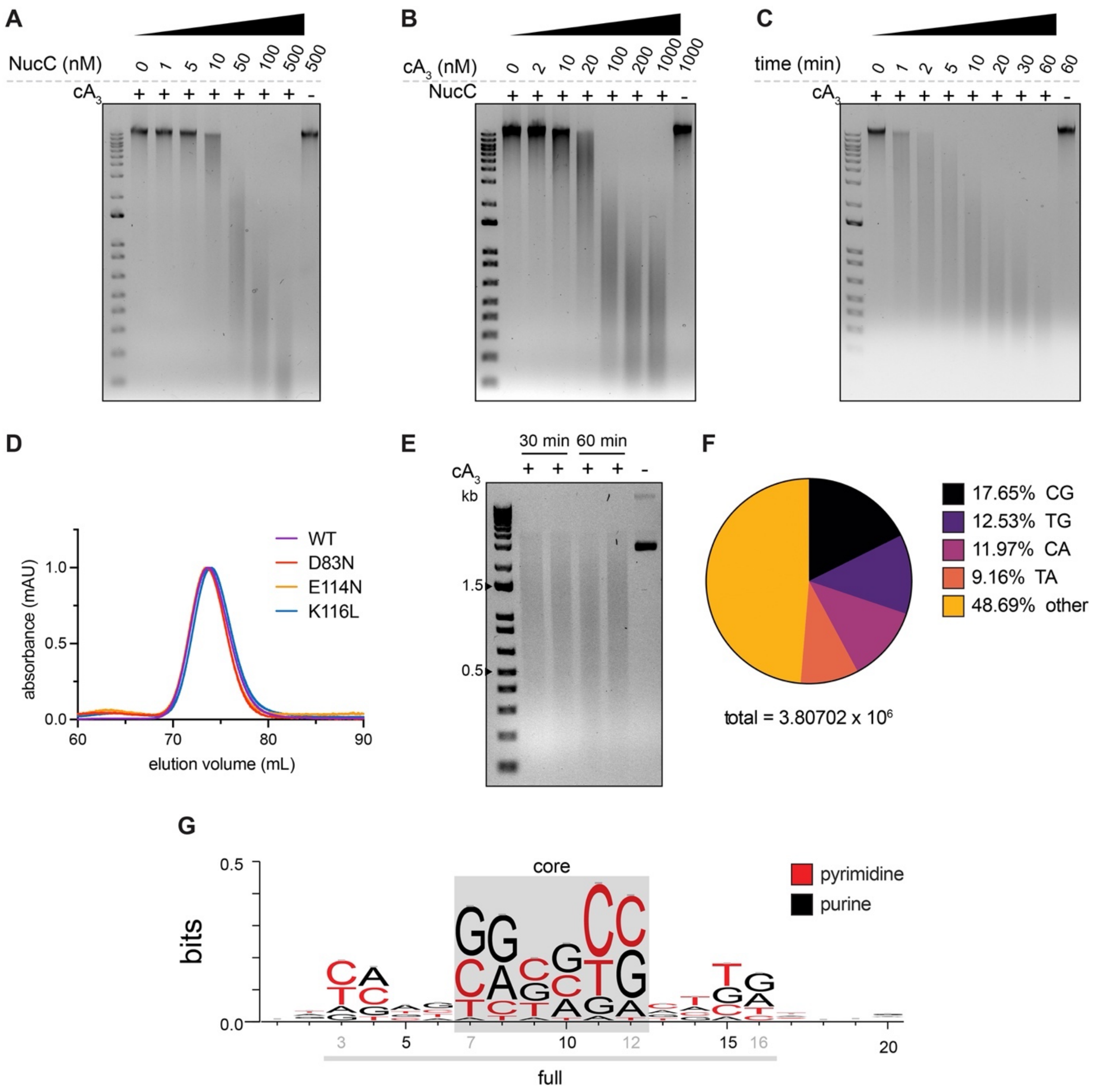


**Supplementary Figure S2. *Serratia* NucC is activated by $cA_3$ and degrades double-stranded DNA *in vitro*.** (**A**) *Serratia* gDNA cleavage assay with increasing concentrations of wild-type NucC in the presence (+, 1000 nM) or absence (-) of $cA_3$. (**B**) *Serratia* gDNA cleavage assay with increasing concentrations of the activator $cA_3$ in the presence (+, 100 nM) or absence (-) of wild-type NucC. (**C**) *Serratia* gDNA cleavage assay with wild-type NucC in the presence of $cA_3$ and increasing incubation times. (**D**) Size exclusion chromatogram of wild-type NucC (purple, pPF2513) and nuclease active site NucC mutants D83N (red, pPF2669), E114N (orange, pPF2671) and K116L (blue, pPF2673). (**E**) NucC cleavage of plasmid for deep sequencing of degradation products. Plasmid pPF1043 was incubated with NucC in the presence of $cA_3$ for 30 min or 60 min. (**F**) Nucleotide preferences at outermost motif positions (3 and 16). Four pyrimidine:purine combinations (C:G, T:G, C:A, T:A) at positions 3 and 16 of the predicted *in vitro* motif account for 51.31% of all sequences. (**G**) Cleavage site preference of NucC, represented as a WebLogo coloured according to pyrimidine (red) or purine (black) nitrogenous base nature. Mapping, per-base coverage and WebLogo summaries are reported in **Supplementary Tables S2**, **S3** and **S4**, respectively.

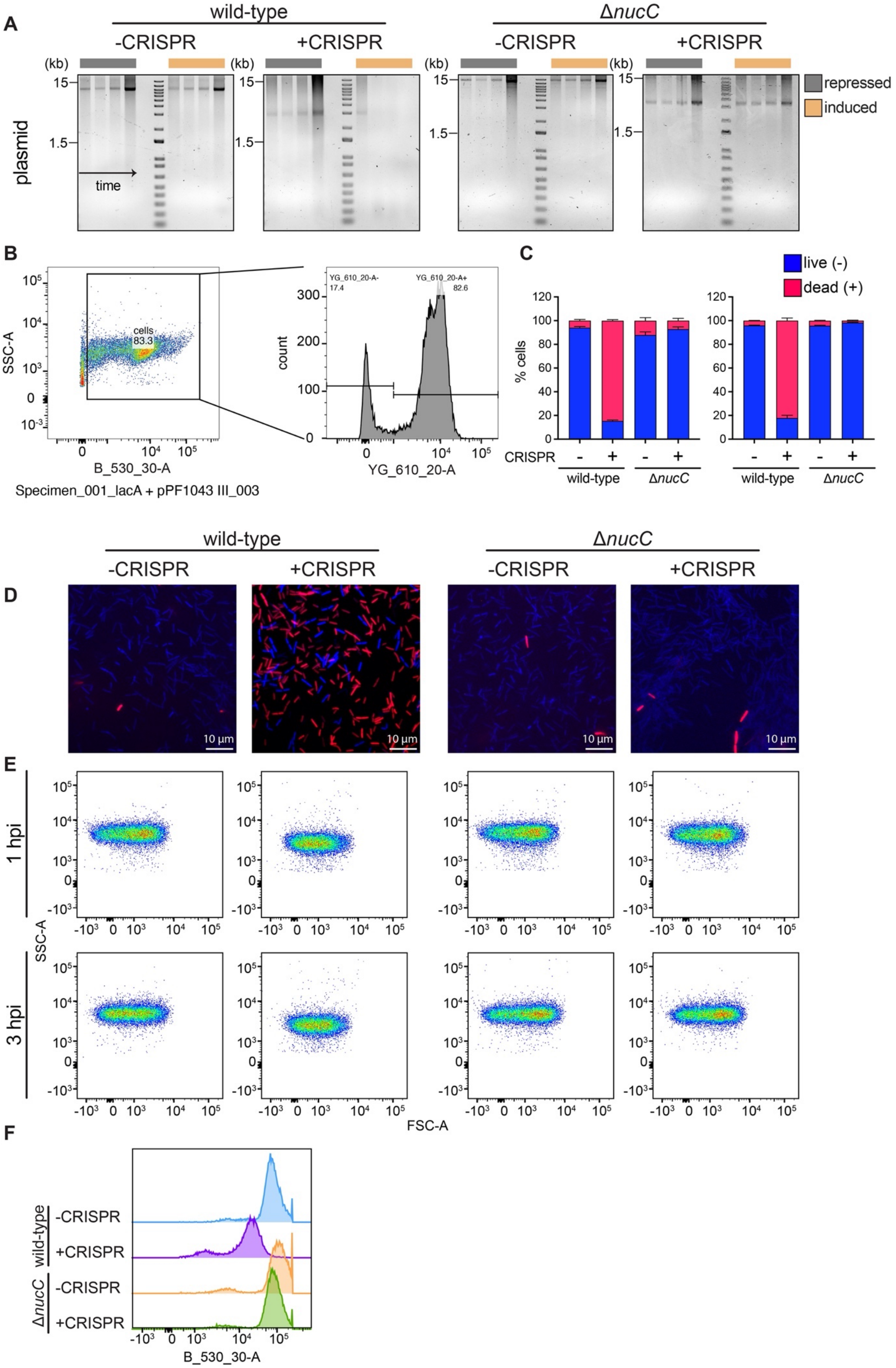
A
wild-type
ΔnucC
-CRISPR
+CRISPR
(kb)
15
1.5
plasmid
time
repressed
induced
B
SSC-A
B_530_30-A
cells
83.3
Specimen_001_lacA + pPF1043 III_003
YG_610_20-A-
17.4
YG_610_20-A+
82.6
count
YG_610_20-A
C
live (-)
dead (+)
% cells
CRISPR
D
10 μm
E
1 hpi
3 hpi
SSC-A
FSC-A
F
B_530_30-A

**Supplementary Figure S3. Type III immunity triggers cell death by destruction of the bacterial genome via NucC.** (**A**) Plasmid DNA was purified from plasmid targeting cultures at 0 h, 0.5 h, 1 h and 3 h post-induction/repression and analysed via gel electrophoresis. (**B**) Flow cytometry gating strategy adopted for cell viability assay during plasmid targeting assay. (**C**) Percentage of live (blue, negative stain) and dead cells (red, positive stain) observed for cell viability assay during plasmid targeting at 1 h and 3 h post-induction (hpi). Cell counts for cell viability assay at 1 hpi and 3 hpi are reported in **Supplementary Table S5**. (**D**) Cell membrane integrity was analysed via confocal microscopy at 3 hpi by staining with propidium iodide which discriminates living cells (blue, negative stain) from dead cells (red, positive stain). (**E**) Forward versus side scatter (FSC vs SSC) of representative samples from cell viability assay during plasmid targeting at 1 hpi and 3 hpi. FSC is indicative of cell size, and SSC relates to the complexity or granularity of the cell. (**F**) SYTO9 fluorescence distributions of representative samples from cell viability assay during plasmid targeting at 1 hpi. The left-skewed distributions of wild-type +CRISPR sample is consistent with DNA degradation.

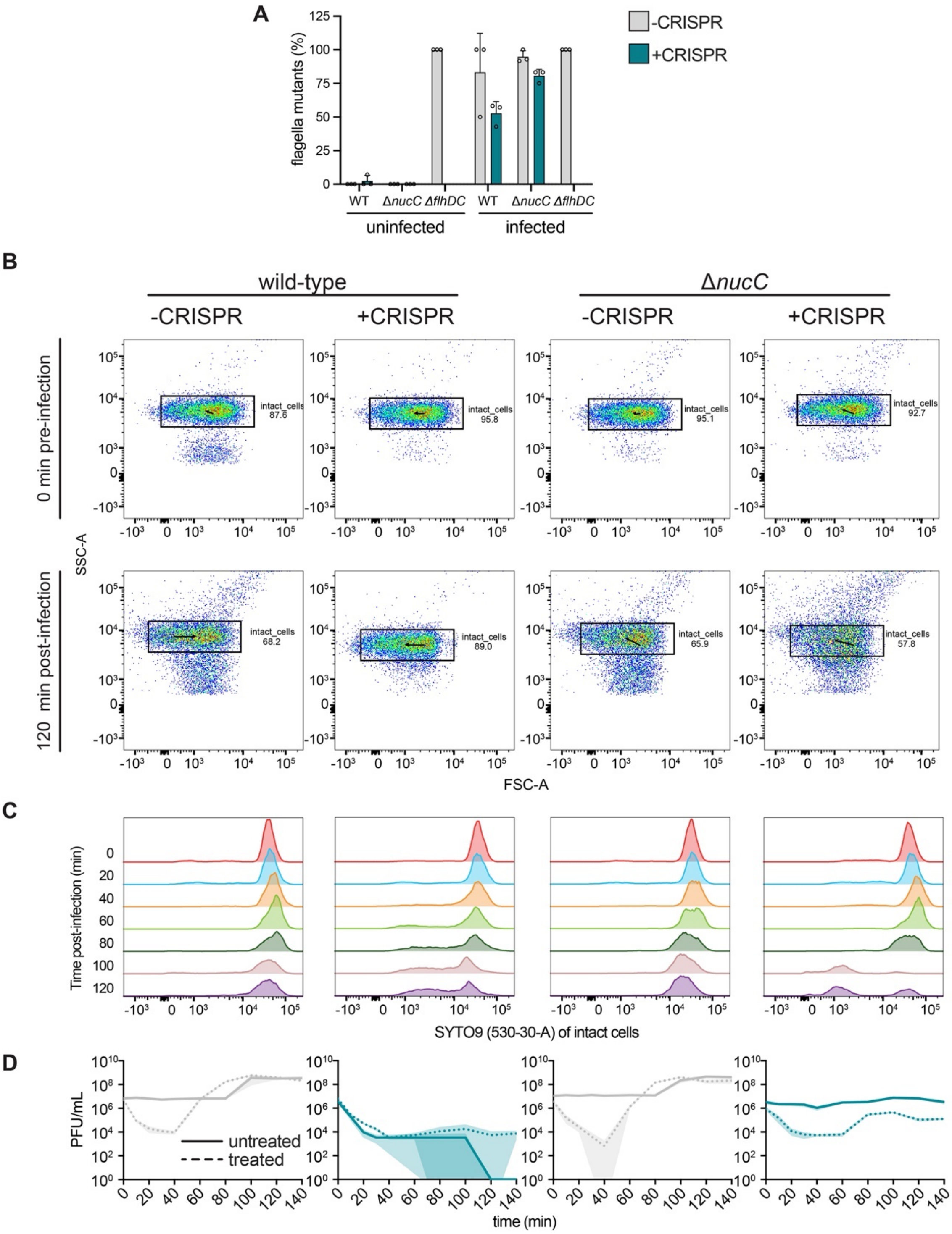


**Supplementary Figure S4. Type III immunity against the jumbo phage results in cell death of the infected individual and prevents jumbo phage reproduction.** (**A**) Percentage of flagella mutants identified after the cell survival assay. A swimming assay was performed from colonies from cell survival assay. (**B**) Scattering profiles of wild-type and *nucC* mutants before and 120 min post-infection. One representative plot of n=3 shown. (**C**) SYTO9 fluorescence distributions of representative samples from (B) and **Figure 4E**. The left-skewed distributions of wild-type anti-phage samples are consistent with DNA degradation. Data for (**B**) and (**C**) shown in **Supplementary Table S6**. (**D**) One-step growth curves performed during jumbo-phage infections and targeting.

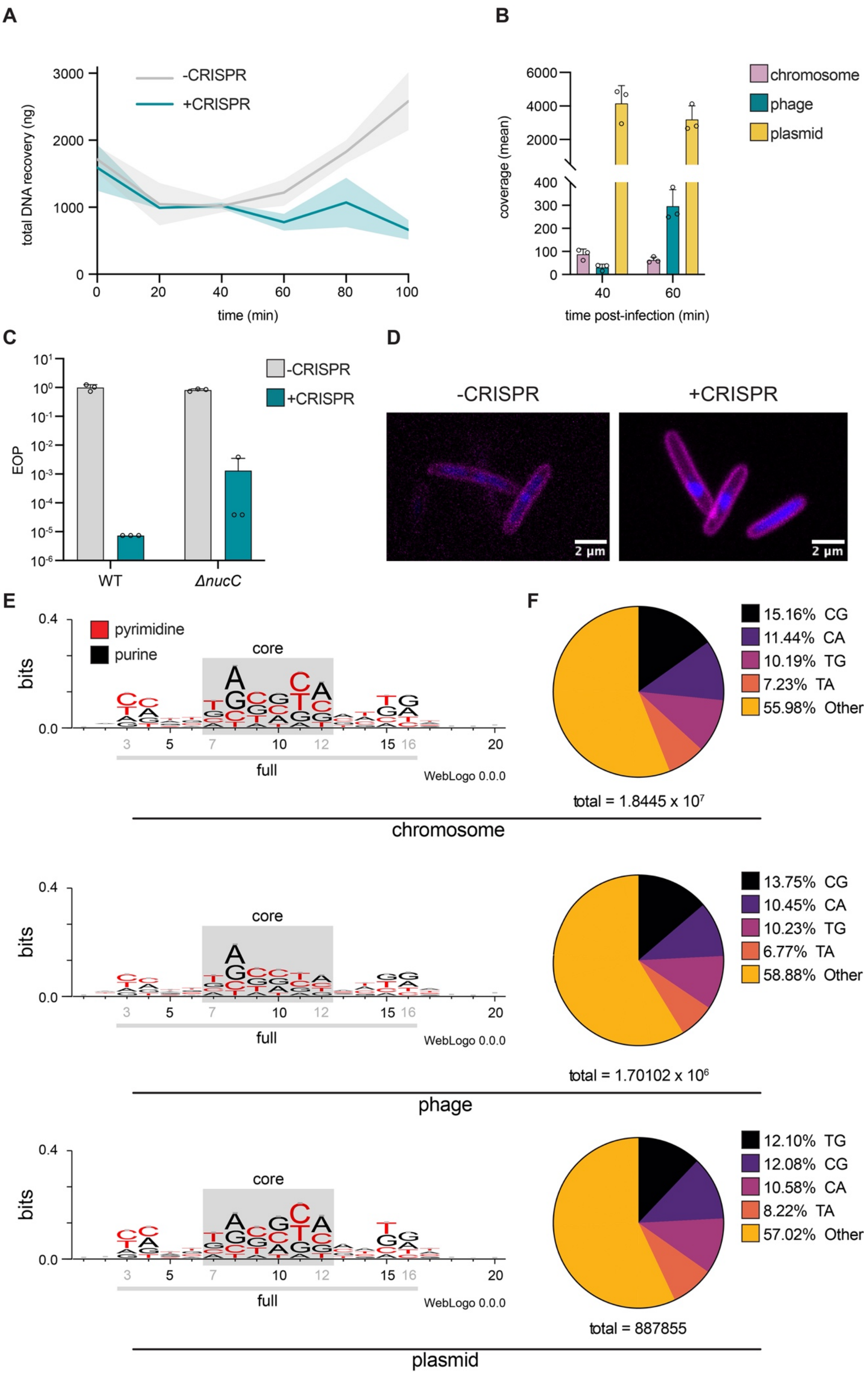

A
B
C
D
E
F
-CRISPR
+CRISPR
total DNA recovery (ng)
time (min)
coverage (mean)
time post-infection (min)
chromosome
phage
plasmid
EOP
WT
ΔnucC
2 µm
pyrimidine
purine
core
full
bits
WebLogo 0.0.0
15.16% CG
11.44% CA
10.19% TG
7.23% TA
55.98% Other
total = 1.8445 x 10^7
13.75% CG
10.45% CA
10.23% TG
6.77% TA
58.88% Other
total = 1.70102 x 10^6
12.10% TG
12.08% CG
10.58% CA
8.22% TA
57.02% Other
total = 887855

**Supplementary Figure S5. The jumbo phage DNA-containing protein shell excludes NucC.** (**A**) Quantification of total DNA extracted throughout a single round of jumbo phage infection from **Figure 5A.** (**B**) Average coverage (per-base) of *Serratia* (chromosome), PCH45 jumbo phage genome (phage) and pPF1467 (plasmid) at 40 and 60 min post-infection. Coverage was calculated using the mosdepth command line tool. The data shown represents biological triplicates plotted as the mean ± standard deviation. (**C**) EOP assay with wild-type (WT) and Δ*nucC* cells expressing mEGFP-NucC with a plasmid with no spacer (-CRISPR) or a spacer targeting PCH45 capsid mRNA (+CRISPR). The data shown represents biological triplicates plotted as the mean ± standard deviation. (**D**) Confocal microscopy of -CRISPR and +CRISPR *Serratia* upon jumbo phage (PCH45) infection. Membranes (magenta) and DNA (blue) were stained with FM4-64 and DAPI, respectively. (**E**) NucC cleaves at partial palindromes in plasmid, phage and chromosomal sequences. Weblogos from deep sequencing of DNA fragments from *in vivo* cleavage by NucC, coloured according to pyrimidine (red) or purine (black) nitrogenous base nature. Cleavage occurs between motif nt positions 10 and 11. (**F**) Nucleotide preferences at outermost motif positions (3 and 16). Four pyrimidine:purine combinations (C:G, T:G, C:A, T:A) at positions 3 and 16 of the predicted *in vivo* motifs account for between 42-51% of all sequences. Mapping, per-base coverage and WebLogo summaries are reported in **Supplementary Tables S2**, **S3** and **S4**, respectively.

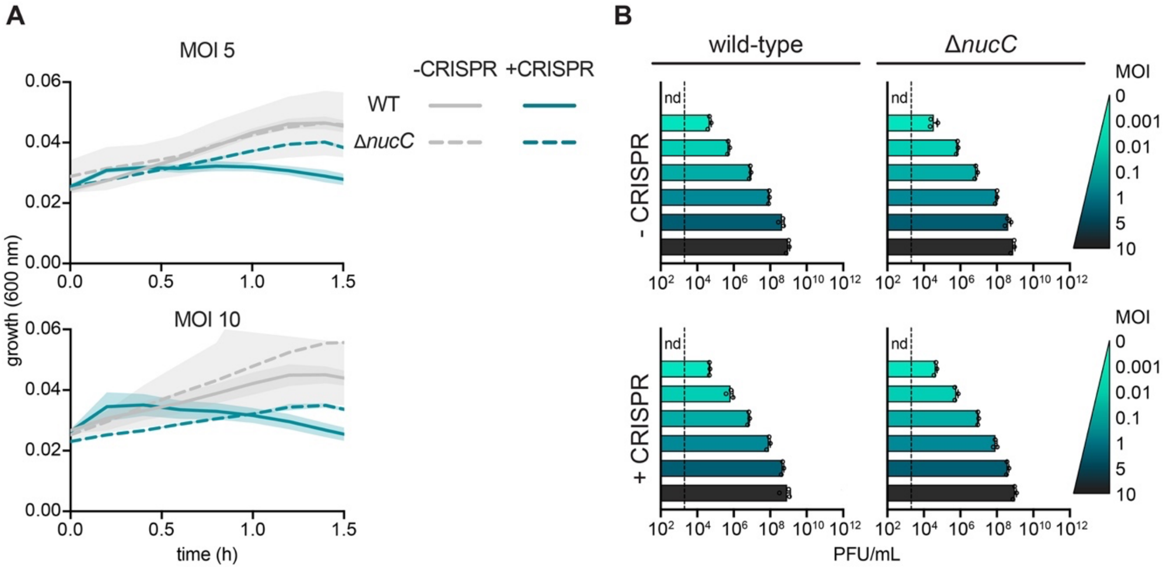


**Supplementary Figure S6. NucC provides jumbo phage immunity by protecting the population at low phage doses.** (**A**) Zoom in first 1.5 h of infection time courses for MOI 5 and MOI 10 from **Figure 6**. (**B**) Initial (0 hpi) phage titres of infection time courses presented in **Figure 6** were measured. Data are biological triplicates plotted as the mean ± standard deviation. Phage infection resulting in no countable plaques were labelled non-determined (nd). Limit of detection of phage titres indicated with a dashed line.

**Supplementary Table S1.** Crystallographic information (**Figure 1**).

| | **Apo-NucC** | **NucC bound to $cA_3$** |
|---|---|---|
| deposition ID | D_1292122076 | D_1292121303 |
| space group | P21 | H32 |
| wavelength | 1.00003 | 1.00003 |
| cell parameters | 79.468 95.492 94.536 | 114.265 114.265 96.069 |
| | 90.000 92.052 90.000 | 90.0 90.0 120.0 |
| resolution | 42.613-1.827 (1.858-1.827) | 43.213-1.480 (1.506-1.480) |
| number of observations | 848140 (41495) | 798761 (38328) |
| number unique | 124878 (6266) | 40117 (1987) |
| Rmerge | 0.049 (0.731) | 0.117 (1.517) |
| Rpim | 0.020 (0.304) | 0.027 (0.354) |
| mean(I)/sd(I) | 23.7 (2.2) | 17.6 (2.2) |
| completeness | 100 (99.9) | 100 (100) |
| multiplicity | 6.8 (6.6) | 19.9 (19.3) |
| CC(1/2) | 1.0 (0.832) | 0.999 (0.773) |
| **refinement** | | |
| reflections used for refinement | 124208 (12390) | 40111 (3970) |
| reflections used for R-free | 6303 (630) | 1962 (192) |
| R-work | 0.1670 (0.2332) | 0.1578 (0.2233) |
| R-free | 0.2076 (0.2848) | 0.1811 (0.2430) |
| RMS(bonds) | 0.01 | 0.011 |
| RMS(angles) | 1.01 | 1.13 |
| ramachandran favored (%) | 97.72 | 96.72 |
| ramachandran outliers (%) | 0.22 | 0 |
| rotamer outliers (%) | 0.83 | 0 |

Statistics for the highest-resolution shell are shown in parentheses.

**Supplementary Table S2.** Mapping summaries resulting from deep sequencing of *in vitro* (**Figure 2**) and *in vivo* (**Figure 5**) degradation products.

| **sample** | **paired-end reads** | **total reads** | **reads aligned to pPF1043** | **reads aligned to PCH45** | **reads aligned to WT** | **reads aligned to pPF1467** | **total reads aligned** | **overall alignment rate (%)** | **unaligned reads** |
|---|---|---|---|---|---|---|---|---|---|
| In_vitro (PE mapping)* | 3884552 | 7769104 | 7292544 | NA | NA | NA | 7292544 | 93.87 | 476560 |
| In_vitro (SE mapping)** | NA | 3884552 | 3808751 | NA | NA | NA | 3808751 | 98.05 | 75801 |
| WT_40_min I | 4660657 | 9321314 | NA | 137370 | 8667662 | 398276 | 9203308 | 98.73 | 118006 |
| WT_40_min II | 4174729 | 8349458 | NA | 131134 | 7740966 | 380078 | 8252178 | 98.83 | 97280 |
| WT_40_min III | 2668582 | 5337164 | NA | 54100 | 4973460 | 237412 | 5264972 | 98.65 | 72192 |
| WT_60_min I | 2847136 | 5694272 | NA | 912266 | 4469758 | 221772 | 5603796 | 98.41 | 90476 |
| WT_60_min II | 4041545 | 8083090 | NA | 1308200 | 6304376 | 340350 | 7952926 | 98.39 | 130164 |
| WT_60_min III | 2958763 | 5917526 | NA | 859610 | 4734120 | 231548 | 5825278 | 98.44 | 92248 |

* this is paired end mapping of full length reads (R1 and R2), used to estimate sequencing depth (coverage) of plasmid

** this is mapping of only R1 (truncated to 15 nt), used for the motif search

**Supplementary Table S3.** Per-base coverage summaries resulting from deep sequencing of *in vitro* (**Figure 2**) and *in vivo* (**Figure 5**) degradation products.

| | **pPF1043** | | | **PCH45** | | | **WT** | | | **pPF1467** | | |
|---|---|---|---|---|---|---|---|---|---|---|---|---|
| **sample** | **mean coverage** | **min coverage** | **max coverage** | **mean coverage** | **min coverage** | **max coverage** | **mean coverage** | **min coverage** | **max coverage** | **mean coverage** | **min coverage** | **max coverage** |
| In_vitro (PE mapping) | 122297.1 | 0 | 343905 | NA | NA | NA | NA | NA | NA | NA | NA | NA |
| WT_40_min I | NA | NA | NA | 39.55 | 0 | 88 | 105.73 | 0 | 680 | 4853.76 | 0 | 9864 |
| WT_40_min II | NA | NA | NA | 38.33 | 0 | 78 | 95.57 | 0 | 715 | 4679.12 | 0 | 9836 |
| WT_40_min III | NA | NA | NA | 15.79 | 0 | 45 | 61.67 | 0 | 416 | 2941.05 | 0 | 6342 |
| WT_60_min I | NA | NA | NA | 262.77 | 0 | 846 | 54.59 | 0 | 1165 | 2667.01 | 0 | 9228 |
| WT_60_min II | NA | NA | NA | 378.26 | 0 | 1246 | 77.47 | 0 | 1570 | 4131.47 | 0 | 16051 |
| WT_60_min III | NA | NA | NA | 249.19 | 0 | 748 | 58.21 | 0 | 1214 | 2805.58 | 0 | 10788 |

**Supplementary Table S4.** Sequence numbers used to build Weblogo motifs from deep sequencing of *in vitro* (**Figure 2**) and *in vivo* (**Figure 5**) degradation products.

| name | total R1 from BAM files | total extracted fasta from BED annotation file | total extracted fasta >20nt | extracted fasta >20nt pPF1043 | extracted fasta >20nt PCH45 | extracted fasta >20nt pPF1467 | extracted fasta >20nt WT |
|---|---|---|---|---|---|---|---|
| In_vitro (R1 mapped) | 3808751 | 3808742 | 3807021 | 3807021 | NA | NA | NA |
| WT_40_min I | 4601654 | 4601454 | 4598153 | NA | 68671 | 195679 | 4333803 |
| WT_40_min II | 4126089 | 4125898 | 4122903 | NA | 65553 | 186891 | 3870459 |
| WT_40_min III | 2632486 | 2632371 | 2630657 | NA | 27047 | 116906 | 2486704 |
| WT_60_min I | 2801898 | 2801670 | 2799256 | NA | 456043 | 108357 | 2234856 |
| WT_60_min II | 3976463 | 3976207 | 3972988 | NA | 653994 | 166830 | 3152164 |
| WT_60_min III | 2912639 | 2912428 | 2909945 | NA | 429717 | 113192 | 2367036 |
| | | | | **total 20nt sequences for building WebLogos** | | | |
| | | | | **pPF1043** | **PCH45** | **pPF1467** | **WT** |
| In_vitro (R1 mapped) | | | | 3807021 | NA | NA | NA |
| WT_concatenation | | | | NA | 1701025 | 887855 | 18445022 |

**Supplementary Table S5.** Flow cytometry viability data during plasmid targeting at 1 hpi and 3 hpi (**Figure 3**).

| | | | | | | | | |
|---|---|---|---|---|---|---|---|---|
| | | | | **1 hour post-induction** | | | | |
| **sample** | **cells \| count** | **cells/PI \| count** | **% dead** | **mean** | **sd** | **% alive** | **mean** | **sd** |
| WT -CRISPR (#1) | 18152 | 1177 | 6.48 | **5.96** | **1.04** | 93.52 | **94.04** | **1.04** |
| WT -CRISPR (#2) | 18747 | 892 | 4.76 | | | 95.24 | | |
| WT -CRISPR (#3) | 18282 | 1214 | 6.64 | | | 93.36 | | |
| WT +CRISPR (#1) | 15548 | 13033 | 83.82 | **84.65** | **0.83** | 16.18 | **15.35** | **0.83** |
| WT +CRISPR (#2) | 16118 | 13641 | 84.63 | | | 15.37 | | |
| WT +CRISPR (#3) | 16094 | 13759 | 85.49 | | | 14.51 | | |
| Δ*nucC* -CRISPR (#1) | 17601 | 2300 | 13.07 | **12.03** | **2.59** | 86.93 | **87.97** | **2.59** |
| Δ*nucC* -CRISPR (#2) | 18317 | 1665 | 9.09 | | | 90.91 | | |
| Δ*nucC* -CRISPR (#3) | 17384 | 2424 | 13.94 | | | 86.06 | | |
| Δ*nucC* +CRISPR (#1) | 18341 | 918 | 5.01 | **7.14** | **1.93** | 94.99 | **92.86** | **1.93** |
| Δ*nucC* +CRISPR (#2) | 18300 | 1404 | 7.67 | | | 92.33 | | |
| Δ*nucC* +CRISPR (#3) | 17984 | 1574 | 8.75 | | | 91.25 | | |
| | | | | **3 hour post-induction** | | | | |
| **sample** | **cells \| count** | **cells/PI \| count** | **% dead** | **mean** | **sd** | **% alive** | **mean** | **sd** |
| WT -CRISPR (#1) | 17623 | 664 | 3.77 | **4.01** | **0.22** | 96.23 | **95.99** | **0.22** |
| WT -CRISPR (#2) | 18424 | 752 | 4.08 | | | 95.92 | | |
| WT -CRISPR (#3) | 19014 | 795 | 4.18 | | | 95.82 | | |
| WT +CRISPR (#1) | 9215 | 7329 | 79.53 | **82.15** | **2.27** | 20.47 | **17.85** | **2.27** |
| WT +CRISPR (#2) | 15970 | 13330 | 83.47 | | | 16.53 | | |
| WT +CRISPR (#3) | 15357 | 12815 | 83.45 | | | 16.55 | | |
| Δ*nucC* -CRISPR (#1) | 19611 | 753 | 3.84 | **4.27** | **0.37** | 96.16 | **95.73** | **0.37** |
| Δ*nucC* -CRISPR (#2) | 18740 | 837 | 4.47 | | | 95.53 | | |
| Δ*nucC* -CRISPR (#3) | 19131 | 862 | 4.51 | | | 95.49 | | |
| Δ*nucC* +CRISPR (#1) | 19549 | 249 | 1.27 | **1.70** | **0.43** | 98.73 | **98.30** | **0.43** |
| Δ*nucC* +CRISPR (#2) | 19172 | 409 | 2.13 | | | 97.87 | | |
| Δ*nucC* +CRISPR (#3) | 19293 | 325 | 1.68 | | | 98.32 | | |

**Supplementary Table S6.** Flow cytometry data during phage infection (**Figure 4**)

| strain | time post-infection (min) | intact_cells \| Count | intact_cells \| Median (B_530_30-A) | intact_cells \| Freq. of Total (10,000) | debris \| Count | debris \| Freq. of Total (10,000) |
|---|---|---|---|---|---|---|
| WT -CRISPR (#1) | 0 | 9085 | 38444 | 90.8 | 915 | 9.2 |
| WT -CRISPR (#2) | 0 | 8115 | 38352 | 81.2 | 1885 | 18.8 |
| WT -CRISPR (#3) | 0 | 8761 | 37984 | 87.6 | 1239 | 12.4 |
| WT -CRISPR (#1) | 20 | 9285 | 40736 | 92.8 | 715 | 7.2 |
| WT -CRISPR (#2) | 20 | 9236 | 40054 | 92.4 | 764 | 7.6 |
| WT -CRISPR (#3) | 20 | 9271 | 40638 | 92.7 | 729 | 7.3 |
| WT -CRISPR (#1) | 40 | 8837 | 50164 | 88.4 | 1163 | 11.6 |
| WT -CRISPR (#2) | 40 | 8734 | 56643 | 87.3 | 1266 | 12.7 |
| WT -CRISPR (#3) | 40 | 8938 | 49921 | 89.4 | 1062 | 10.6 |
| WT -CRISPR (#1) | 60 | 8329 | 56643 | 83.3 | 1671 | 16.7 |
| WT -CRISPR (#2) | 60 | 7933 | 60050 | 79.3 | 2067 | 20.7 |
| WT -CRISPR (#3) | 60 | 7731 | 63510 | 77.3 | 2269 | 22.7 |
| WT -CRISPR (#1) | 80 | 6439 | 51272 | 64.4 | 3561 | 35.6 |
| WT -CRISPR (#2) | 80 | 6855 | 52278 | 68.5 | 3145 | 31.5 |
| WT -CRISPR (#3) | 80 | 6862 | 44226 | 68.6 | 3138 | 31.4 |
| WT -CRISPR (#1) | 100 | 6083 | 32486 | 60.8 | 3917 | 39.2 |
| WT -CRISPR (#2) | 100 | 3067 | 27085 | 30.7 | 6933 | 69.3 |
| WT -CRISPR (#3) | 100 | 5805 | 30232 | 58.1 | 4195 | 41.9 |
| WT -CRISPR (#1) | 120 | 6710 | 33516 | 67.1 | 3290 | 32.9 |
| WT -CRISPR (#2) | 120 | 7084 | 29517 | 70.8 | 2916 | 29.2 |
| WT -CRISPR (#3) | 120 | 6846 | 30016 | 68.5 | 3154 | 31.5 |
| WT +CRISPR (#1) | 0 | 9293 | 34914 | 92.9 | 707 | 7.1 |
| WT +CRISPR (#2) | 0 | 9176 | 38259 | 91.8 | 824 | 8.2 |
| WT +CRISPR (#3) | 0 | 9577 | 33597 | 95.8 | 423 | 4.2 |
| WT +CRISPR (#1) | 20 | 9314 | 33759 | 93.1 | 686 | 6.9 |
| WT +CRISPR (#2) | 20 | 9265 | 39193 | 92.7 | 735 | 7.3 |
| WT +CRISPR (#3) | 20 | 9443 | 33840 | 94.4 | 557 | 5.6 |
| WT +CRISPR (#1) | 40 | 9213 | 31263 | 92.1 | 787 | 7.9 |
| WT +CRISPR (#2) | 40 | 9108 | 29873 | 91.1 | 892 | 8.9 |
| WT +CRISPR (#3) | 40 | 9262 | 32253 | 92.6 | 738 | 7.4 |
| WT +CRISPR (#1) | 60 | 8968 | 25038 | 89.7 | 1032 | 10.3 |
| WT +CRISPR (#2) | 60 | 9054 | 25458 | 90.5 | 946 | 9.5 |
| WT +CRISPR (#3) | 60 | 9089 | 22079 | 90.9 | 911 | 9.1 |
| WT +CRISPR (#1) | 80 | 8831 | 20472 | 88.3 | 1169 | 11.7 |
| WT +CRISPR (#2) | 80 | 8885 | 19438 | 88.8 | 1115 | 11.2 |
| WT +CRISPR (#3) | 80 | 8956 | 20472 | 89.6 | 1044 | 10.4 |
| WT +CRISPR (#1) | 100 | 9083 | 10650 | 90.8 | 917 | 9.2 |
| WT +CRISPR (#2) | 100 | 8913 | 6502 | 89.1 | 1087 | 10.9 |
| WT +CRISPR (#3) | 100 | 9019 | 9884 | 90.2 | 981 | 9.8 |
| WT +CRISPR (#1) | 120 | 8695 | 8281 | 87 | 1305 | 13.0 |
| WT +CRISPR (#2) | 120 | 8486 | 6253 | 84.9 | 1514 | 15.1 |

| | | | | | | |
|---|---|---|---|---|---|---|
| WT +CRISPR (#3) | 120 | 8898 | 10554 | 89 | 1102 | 11.0 |
| Δ*nucC* -CRISPR (#1) | 0 | 9588 | 30670 | 95.9 | 412 | 4.1 |
| Δ*nucC* -CRISPR (#2) | 0 | 9535 | 34580 | 95.3 | 465 | 4.7 |
| Δ*nucC* -CRISPR (#3) | 0 | 9512 | 33037 | 95.1 | 488 | 4.9 |
| Δ*nucC* -CRISPR (#1) | 20 | 9250 | 31869 | 92.5 | 750 | 7.5 |
| Δ*nucC* -CRISPR (#2) | 20 | 9315 | 37258 | 93.2 | 685 | 6.8 |
| Δ*nucC* -CRISPR (#3) | 20 | 9265 | 36372 | 92.7 | 735 | 7.3 |
| Δ*nucC* -CRISPR (#1) | 40 | 8811 | 36812 | 88.1 | 1189 | 11.9 |
| Δ*nucC* -CRISPR (#2) | 40 | 8662 | 46420 | 86.6 | 1338 | 13.4 |
| Δ*nucC* -CRISPR (#3) | 40 | 8771 | 45972 | 87.7 | 1229 | 12.3 |
| Δ*nucC* -CRISPR (#1) | 60 | 8352 | 34085 | 83.5 | 1648 | 16.5 |
| Δ*nucC* -CRISPR (#2) | 60 | 8025 | 46985 | 80.2 | 1975 | 19.8 |
| Δ*nucC* -CRISPR (#3) | 60 | 8633 | 41531 | 86.3 | 1367 | 13.7 |
| Δ*nucC* -CRISPR (#1) | 80 | 8011 | 24450 | 80.1 | 1989 | 19.9 |
| Δ*nucC* -CRISPR (#2) | 80 | 7360 | 33921 | 73.6 | 2640 | 26.4 |
| Δ*nucC* -CRISPR (#3) | 80 | 8106 | 29944 | 81.1 | 1894 | 18.9 |
| Δ*nucC* -CRISPR (#1) | 100 | 8160 | 20617 | 81.6 | 1840 | 18.4 |
| Δ*nucC* -CRISPR (#2) | 100 | 6837 | 26764 | 68.4 | 3163 | 31.6 |
| Δ*nucC* -CRISPR (#3) | 100 | 7770 | 24218 | 77.7 | 2230 | 22.3 |
| Δ*nucC* -CRISPR (#1) | 120 | 8368 | 19995 | 82.7 | 1632 | 17.3 |
| Δ*nucC* -CRISPR (#2) | 120 | 7315 | 25458 | 73.2 | 2685 | 26.8 |
| Δ*nucC* -CRISPR (#3) | 120 | 6588 | 25702 | 65.9 | 3412 | 34.1 |
| Δ*nucC* +CRISPR (#1) | 0 | 9343 | 38352 | 93.4 | 657 | 6.6 |
| Δ*nucC* +CRISPR (#2) | 0 | 9460 | 34085 | 94.6 | 540 | 5.4 |
| Δ*nucC* +CRISPR (#3) | 0 | 9265 | 38352 | 92.7 | 735 | 7.3 |
| Δ*nucC* +CRISPR (#1) | 20 | 9178 | 41732 | 91.8 | 822 | 8.2 |
| Δ*nucC* +CRISPR (#2) | 20 | 9176 | 38630 | 91.8 | 824 | 8.2 |
| Δ*nucC* +CRISPR (#3) | 20 | 9185 | 41132 | 91.8 | 815 | 8.2 |
| Δ*nucC* +CRISPR (#1) | 40 | 8665 | 58463 | 86.7 | 1335 | 13.3 |
| Δ*nucC* +CRISPR (#2) | 40 | 8795 | 52405 | 87.9 | 1205 | 12.1 |
| Δ*nucC* +CRISPR (#3) | 40 | 8611 | 61982 | 86.1 | 1389 | 13.9 |
| Δ*nucC* +CRISPR (#1) | 60 | 8002 | 61530 | 80 | 1998 | 20.0 |
| Δ*nucC* +CRISPR (#2) | 60 | 8497 | 50900 | 85 | 1503 | 15.0 |
| Δ*nucC* +CRISPR (#3) | 60 | 8027 | 59179 | 80.3 | 1973 | 19.7 |
| Δ*nucC* +CRISPR (#1) | 80 | 7387 | 34331 | 73.9 | 2613 | 26.1 |
| Δ*nucC* +CRISPR (#2) | 80 | 5624 | 43694 | 56.2 | 4376 | 43.8 |
| Δ*nucC* +CRISPR (#3) | 80 | 5547 | 44119 | 55.5 | 4453 | 44.5 |
| Δ*nucC* +CRISPR (#1) | 100 | 3664 | 1407 | 36.6 | 6336 | 63.4 |
| Δ*nucC* +CRISPR (#2) | 100 | 5809 | 21820 | 58.1 | 4191 | 41.9 |
| Δ*nucC* +CRISPR (#3) | 100 | 4910 | 1034 | 49.1 | 5090 | 50.9 |
| Δ*nucC* +CRISPR (#1) | 120 | 5637 | 1804 | 56.4 | 4363 | 43.6 |
| Δ*nucC* +CRISPR (#2) | 120 | 6276 | 21975 | 62.8 | 3724 | 37.2 |
| Δ*nucC* +CRISPR (#3) | 120 | 5781 | 16847 | 57.8 | 4219 | 42.2 |

**Supplementary Table S7. Strains used in this study.**

| name | genotype/phenotype | reference |
|---|---|---|
| ***Serratia* sp ATCC 39006** | | |
| LacA | Lac- EMS mutant (“wild-type” parental strain for this study) | (Thomson et al., 2000) |
| PCF686 | LacA-derivative, Δ*nucC* | (Malone et al., 2020) |
| PCF879 | LacA-derivative, Δ*flhDC*::Cm | (Hampton et al., 2016) |
| ***Escherichia coli*** | | |
| DH5α | cloning strain. $F^-$ , φ80Δd*lac*ZM15, Δ(*lacZYA*–arg*F*)U169, *endA1*, *recA1*, *hsdR*17 ($rK^-$ $mK^+$ ), *deoR*, *thi-1*, *sup*E44, $λ^-$, *gyr*A96, *rel*A1 | (Taylor et al., 1993) |
| ST18 | auxotrophic donor for biparental conjugation. S17-1 λ*pir* Δ*hemA* | (Jackson et al., 2020; Thoma and Schobert, 2009) |
| BL21(DE3) | protein expression strain. Str., B, $F^–$, *ompT*, *gal*, *dcm*, *lon*, $hsdS_B(r_B^- m_B^-)$, λ(DE3, [*lacI*, *lacUV5-T7p07*, *ind1*, *sam7*, *nin5*]), $[malB^+]_{K-12}(λ^S)$ | (Studier and Moffatt, 1986) |
| **Bacteriophages** | | |
| PCH45 | lytic jumbo phage, family *Myoviridae;* infects *Serratia* sp. ATCC 39006 | (Malone *et al.*, 2020) |
| JS26 | lytic non-jumbo phage, family *Siphoviridae*; infects Serratia sp. ATCC 39006 | (Malone et al., 2022) |

**Supplementary Table S8. Oligonucleotides used in this study.**

| name | sequence (5′-3′) | description |
|---|---|---|
| **cloning** | | |
| pPF2513-F | TACTTCCAATCCAATGCAATGACTAATCAGGCAAAAAA | F primer for cloning *Serratia nucC* into expression vector, generating pPF2513 |
| pPF2513-R | TTATCCACTTCCAATGTTATTATTCCAGACTATCTATAT | R primer for cloning *Serratia nucC* into expression vector, generating pPF2513 |
| PF2767 | AGACTCAGGTCGTCTTTGTGCGCTTTGTAGAAAGCAGCG | F primer complement of PF2768 to generate spacer targeting phage JS26 in pPF1477 |
| PF2768 | AGGACGCTGCTTTCTACAAAGCGCACAAAGACGACCTGA | R primer complement of PF2767 to generate spacer targeting phage JS26 in pPF1477 |
| PF3896 | CAAAGAGGAGAAATTAACTATGACTAATCAGGCAAAAAAGTTATCTA G | F1 primer for Gibson assembly of *Serratia nucC* into pQE-80L-stuffer, generating pPF2007 |
| PF3897 | TCATACTAGGATCCGCATGCAAAGAGGAGAAATTAACTATGACTAAT CAGG | F2 primer for Gibson assembly of *Serratia nucC* into pQE-80L-stuffer, generating pPF2007 |
| PF3898 | TGGCTGCAGGTCGACCCGGGTTATTCCAGACTATCTATATACACCC GC | R primer for Gibson assembly of *Serratia nucC* into pQE-80L-stuffer, generating pPF2007 |
| PF4688 | GCGAATTCGAGCTCGGTACCAAAGAGGAGAAATTAACTATGGTGAG | F primer for amplification of gBlock PF3809, overlap with pBAD30 for Gibson assembly (KpnI*);* pPF2290 cloning |
| PF4689 | CTTTTTTGCCTGATTAGTCATGGATCCGCCTCCACCG | R primer for amplification of gBlock PF3809, overlap with PF4690; pPF2290 cloning |
| PF4690 | AGGGCGGTGGAGGCGGATCCATGACTAATCAGGCAAAAAAGTTATC | F primer for amplification of *Serratia nucC* + linker (Glyx5-Ser), overlap with PF4689; pPF2290 cloning |
| PF4691 | CAAAAGGTCATCCACTGCAGTTATTCCAGACTATCTATATACACCC | R primer for amplification of *Serratia nucC*, overlap with pBAD30 for Gibson assembly (PstI*)*; pPF2290 cloning |
| PF3809 | TCGTCTTCACCTCGAGAAATCAAAGAGGAGAAATTAACTATGGTGAG CAAGGGCGAGGAGCTGTTCACCGGGGTGGTGCCCATCCTGGTCGA GCTGGACGGCGACGTAAACGGCCACAAGTTCAGCGTGTCCGGCGA GGGCGAGGGCGATGCCACCTACGGCAAGCTGACCCTGAAGTTCAT CTGCACCACCGGCAAGCTGCCCGTGCCCTGGCCCACCCTCGTGAC CACCCTGACCTACGGCGTGCAGTGCTTCAGCCGCTACCCCGACCA CATGAAGCAGCACGACTTCTTCAAGTCCGCCATGCCCGAAGGCTAC GTCCAGGAGCGCACCATCTTCTTCAAGGACGACGGCAACTACAAGA CCCGCGCCGAGGTGAAGTTCGAGGGCGACACCCTGGTGAACCGCA TCGAGCTGAAGGGCATCGACTTCAAGGAGGACGGCAACATCCTGG GGCACAAGCTGGAGTACAACTACAACAGCCACAACGTCTATATCAT GGCCGACAAGCAGAAGAACGGCATCAAGGTGAACTTCAAGATCCGC CACAACATCGAGGACGGCAGCGTGCAGCTCGCCGACCACTACCAG CAGAACACCCCCATCGGCGACGGCCCCGTGCTGCTGCCCGACAAC CACTACCTGAGCACCCAGTCCAAGCTGAGCAAAGACCCCAACGAGA AGCGCGATCACATGGTCCTGCTGGAGTTCGTGACCGCCGCCGGGA TCACTCTCGGCATGGACGAGCTGTACAAGGGCGGTGGAGGCGGAT CCCCTGTTGATAGATCCAGTAATGAC | gBlock template for amplification of RBS-mEGFP(no STOP codon)-linker(Gly5x-Ser); pPF2290 cloning |
| PF5145 | TATAGAATTCAAAGAGGAGAAATTAACTATGACTAATCAGGCAAAAA AGT | F primer for cloning *Serratia nucC* + artificial RBS into pPF1618, generating pPF2503 (EcoRI) |
| PF5146 | TATAAAGCTTTTATTCCAGACTATCTATATACACCCGCC | R primer for cloning *Serratia nucC* + artificial RBS into pPF1618, generating pPF2503 (HindIII) |

| name | sequence (5′-3′) | description |
|---|---|---|
| PF5149 | TATAGAATTCAAAGAGGAGAAATTAACTATGGAGGACCTTGATGCG | F primer for cloning *Thermus thermophilus csm6* + artificial RBS into pPF1618, generating pPF2505 (EcoRI) |
| PF5150 | TATAAAGCTTTTAGAACCCCAAGGGTACGGGT | R primer for cloning *Thermus thermophilus csm6* + artificial RBS into pPF1618, generating pPF2505 (HindIII) |
| PF5539 | CCAGATAAATGCAGTGATTTTTG | F primer for site-directed mutagenesis of *Serratia nucC* active site (D83N) in pPF2513, generating pPF2669 |
| PF5540 | TGCATTTATCTGGTCGCTG | R primer for site-directed mutagenesis of Serratia *nucC* active site (D83N) in pPF2513, generating pPF2669 |
| PF5541 | GTACTGAATGTTAAACCAACCAT | F primer for site-directed mutagenesis of Serratia *nucC* active site (E114N) in pPF2513, generating pPF2671 |
| PF5542 | TTAACATTCAGTACCGCGTAC | R primer for site-directed mutagenesis of Serratia *nucC* active site (E114N) in pPF2513, generating pPF2671 |
| PF5543 | GGTTCTTCCAACCATTAATAAAACC | F primer for site-directed mutagenesis of Serratia *nucC* active site (K116L) in pPF2513, generating pPF2673 |
| PF5544 | GTTGGAAGAACCTCCAGTA | R primer for site-directed mutagenesis of Serratia *nucC* active site (K116L) in pPF2513, generating pPF2673 |
| **NucC cleavage assays (Figure 2G-H)** | | |
| PF73 | GACTCTAGACACGTGGAGAAACCAAAGCC | F primer for amplification of a *Serratia* chromosomic region, generating a 1419 bp product for cleavage assays. Binds XRE family transcriptional regulator CDS |
| PF807 | GATCCCGGGTCAGTTCCTTGCCGTAGC | R primer for amplification of a *Serratia* chromosomic region, generating a 1419 bp product for cleavage assays. Binds DUF165 domain-containing protein CDS |
| PF6283 | CCCTACGCTCCCTCCAGCGCTGTCGGGGATATAGTCACTCGGAGTT AGAGAGTTTTAGGATTGATTACTGAACTCTAGTATGGTAAACTGTGA AAACTCATAAAGCTGACGAAGTAAAAGAATCAAACTAATAACTCAAT CCAGTCTAAAGAGTAGAAAGTTGGTGAAAGATTGTGAGTCAGTCACT TAATGGTCTTAGA | no motif negative control gBlock |
| PF6284 | CCCTACGCTCCCTCCAGCGCTGTCGGGGATATAGTCACTCGGCAAG GGCGCCCTTGAGGATTGATTACTGAACTCTAGTATGGTAAACTGTGA AAACTCATAAAGCTGACGAAGTAAAAGAATCAAACTAATAACTCAAT CCAGTCTAAAGAGTAGAAAGTTGGTGAAAGATTGTGAGTCAGTCACT TAATGGTCTTAGA | full motif gBlock |
| PF6285 | CCCTACGCTCCCTCCAGCGCTGTCGGGGATATAGTCACTCGGAGTT GGCGCCTTTTAGGATTGATTACTGAACTCTAGTATGGTAAACTGTGA AAACTCATAAAGCTGACGAAGTAAAAGAATCAAACTAATAACTCAAT CCAGTCTAAAGAGTAGAAAGTTGGTGAAAGATTGTGAGTCAGTCACT TAATGGTCTTAGA | core motif gBlock |
| **screening** | | |
| PF2202 | TATTGCATGCGGCTGACGATCTGGCGTC | chromosomal *Serratia nucC*, F primer |
| PF2199 | TCTTGGATCCGCTAGCGGCCTGCCGGAGAAC | chromosomal *Serratia nucC*, R primer |
| PF138 | CACACTTTGCTATGCCATAG | pPF781-derived plasmids, F primer |
| PF1702 | CGAAGACGAAAGGGCCTCGTGATACGCAAGCTTTATGGCTTGTAAA CCGTTTTGTG | pPF781-derived plasmids, R primer |

| name | sequence (5′-3′) | description |
|---|---|---|
| PF4181 | AAAGAAATCATAAAAAATTTATTTGCTTTGTGAGCGGAT | pPF976-derived plasmids, F primer |
| PF3737 | TTTATGCATCTTCAGTCAGGGAGCGTC | pPF976-derived plasmids, R primer |
| PF2231 | TTTTACTAGTAGACGTTCAACAACGTCATG | PCH45 capsid gene, F primer |
| PF2232 | TTTTGGTACCGAAGTTATATTCGCGCGGTG | PCH45 capsid gene, R primer |
| PF138 | CACACTTTGCTATGCCATAG | pPF1618-derived plasmids, F primer |
| PF210 | GTCATTACTGGATCTATCAACAGG | pPF1618-derived plasmids, R primer |

**Supplementary Table S9. Plasmids used in this study.**

| name | description | features | construction | reference |
|---|---|---|---|---|
| **protein expression and purification (Figs. 1 and 2)** | | | | |
| pPF2007 | template for *nucC* cloning into expression vector | pBR322/ori, RP4/oriT, ApR, lacI/T5 | Gibson assembly using PF3896, PF3897 + PF3898 and SphI and SmaI to Gibson assemble *nucC* into pQE-80L-oriT stuffer | This study |
| pPF2513 | 6xHis-TEV-NucC expression vector | ColE1/ori, f1/ori, KmR, lacI/T7 | pPF2513-F + pPF2513-R paired in a PCR to amplify *nucC* from pPF2007 and clone it into an expression vector through Ligation Independent Cloning. | This study |
| pPF2669 | 6xHis-TEV-NucC D83N mutant expression vector | ColE1/ori, f1/ori, KmR, lacI/T7 | PF5539 + PF5540 paired in a PCR to introduce NucC D83N mutation in pPF2513 | This study |
| pPF2671 | 6xHis-TEV-NucC E114N mutant expression vector | ColE1/ori, f1/ori, KmR, lacI/T7 | PF5541 + PF5542 paired in a PCR to introduce NucC E114N mutation in pPF2513 | This study |
| pPF2673 | 6xHis-TEV-NucC K116L mutant expression vector | ColE1/ori, f1/ori, KmR, lacI/T7 | PF5543 + PF5544 paired in a PCR to introduce NucC K116L mutation in pPF2513 | This study |
| **plasmid targeting (Fig. 3)** | | | | |
| pPF781 | untargeted control for the type III-A system | p15A/ori, RP4/oriT, CmR, pBAD/araC | pBAD30 derivative | (Patterson et al., 2016) |
| pPF1043 | targeted type III-A with protospacer complementary to *Serratia* CRISPR3 spacer 1 | p15A/ori, RP4/oriT, CmR, pBAD/araC | pPF781 derivative | (Patterson *et al.*, 2016) |
| **phage targeting (Figs. 4, 5, 6 and 7)** | | | | |
| pPF976 | type III-A repeat-BsaI-repeat construct for artificial crRNA | pBR322/ori, RP4/oriT, KmR, lacI/T5 | pPF260 derivative | (Malone *et al.*, 2020) |
| pPF1467 | anti-PCH45 III-A spacer overexpression. III-A_PCH45_PS4 (capsid protein) | pBR322/ori, RP4/oriT, KmR, lacI/T5 | pPF976 derivative | (Malone *et al.*, 2020) |
| pPF1477 | anti-JS26 III-A spacer overexpression. III-A_JS26_PS8 (capsid protein) | pBR322/ori, RP4/oriT, KmR, lacI/T5 | PF2767 and PF2768 were annealed, digested with BsaI and ligated into pPF976 cut with BsaI | This study |
| **NucC localization microscopy (Fig. 5)** | | | | |
| pPF2290 | mEGFP-NucC expression vector | p15A/ori, RP4/oriT, GmR, pBAD/araC | Gibson assembly PF4688/PF4689 (gblockPF3809) + PF4690/PF4691 (LacA) | This study |
| **accessory nuclease swap (Fig. 7)** | | | | |
| pPF1618 | empty vector control for expression | RK2/ori, OriT, ApR, pBAD/araC | pSEVA1810 (pBAD) and pSEVA121 (ApR, RK2) were digested with PacI and EcoRI, and ligated into one vector. | This study |
| pPF2503 | *Serratia* NucC expression vector | RK2/ori, OriT, ApR, pBAD/araC | PF5145 + PF5146 paired in a PCR to amplify *nucC* from *Serratia* LacA and cloned it into pPF1618 (EcoRI/HindIII) | This study |
| pC0075 | *Thermus thermophilus* Csm6 template | | TtCsm6 His6-TwinStrep-SUMO-BsaI (Addgene #115270) | (Gootenberg et al., 2018) |
| pPF2505 | *T. thermophilus* Csm6 expression vector | RK2/ori, OriT, ApR, pBAD/araC | PF5149 + PF5150 paired in a PCR to amplify *csm6* from pAddgene_115270 and clone it into pPF1618 (EcoRI/HindIII) | This study |
